# Novel populations of Tr1 cells contribute to the resolution of acute influenza A virus infection

**DOI:** 10.1101/2022.12.06.519368

**Authors:** Caitlin A Abbott, Emily L Freimayer, Timona S Tyllis, Todd S Norton, Mohammed Alsharifi, Aaron H S Heng, Stephen M Pederson, Zhipeng Qu, Mark Armstrong, Geoffrey R Hill, Shaun R McColl, Iain Comerford

**Author notes:** Equal contributions (Shaun R McColl, Iain Comerford).

## Abstract

Type I regulatory (Tr1) cells contribute to immune suppression in the context of chronic infection, autoimmunity, and transplant tolerance. However, their physiological relevance in the resolution of acute respiratory infection is not understood. Here, we identify Tr1 cells accumulating in the lung parenchyma during resolution of the response to sublethal influenza A virus infection in mice. Tr1 cells were dependent on IL-27Rα and in their absence recovery from IAV-induced weight loss is impaired. Notably, these Tr1 cells did not necessarily co-express the typical Tr1 markers LAG-3 and CD49b, with four distinct populations of Tr1 cells apparent in the lungs. Each population was suppressive and were differentially dependent on IL-10 to mediate suppression. Transcriptional analysis revealed a core Tr1 gene signature in each population and distinct expression profiles indicative of different states of activation and differentiation. Finally, sort-transfer experiments indicated non-linear plasticity between these subsets of Tr1 cells. Together, these data support Tr1 cells contributing to the resolution of acute inflammation and define novel Tr1 cell phenotypes in acute infection.

## Introduction

Respiratory infections caused by pathogens such as influenza A virus (IAV) and SARS-CoV2 are a cause of significant morbidity and mortality worldwide. Understanding how immune activation and suppression are balanced during acute respiratory infection is critical for developing effective therapeutic interventions to promote tissue recovery and pathogen clearance. Antigen-specific immune suppression mediated by regulatory T cells (Tregs) plays an important role in control of the immune response, both in terms of its magnitude and resolution. FOXP3^+^ Tregs are the best characterised T cell population involved in regulation of the adaptive immune response to IAV, having been shown to permit establishment of appropriate protective responses (Ruckwardt et al., 2009; Sehrawat et al., 2008), constrain pathological effector T cell responses (Rogers et al., 2018; Lund et al., 2008; Fulton et al., 2010; Zou et al., 2014; Lu et al., 2019), and promote tissue repair (Arpaia et al., 2015). However, other regulatory T cell subsets may also make important contributions during acute infection.

Tr1 cells were first described as IL-10-secreting CD4^+^ T cells with a suppressive function that often co-express the Th1 cytokine IFNγ (Roncarolo et al., 1988; Bacchetta et al., 1990; Groux et al., 1997; Heinzel et al., 1991; Ghalib et al., 1993; Holaday et al., 1993; Reiner et al., 1994; Reed et al., 1994; Hagenbaugh et al., 1997). However, current classification of this subset has been refined to exclude other subsets of CD4^+^ T cells that have the capacity to secrete IL-10, such as Tregs, Th2, Th3, Th9, Th17, and other T cells that lack suppressive functionality. Specifically, Tr1 cells are now generally accepted to be CD4^+^ T cells that do not express FOXP3 or produce cytokines associated with Th2 or Th17 cells and that suppress bystander T cell activation (Roncarolo et al., 2018; Bacchetta et al., 1990; Groux et al., 1997). These cells develop in an IL-27-dependent manner from naïve precursors *in vitro* (Groux et al., 1997; Pot et al., 2009; Batten et al., 2008; Zhang et al., 2020) and can also emerge from other T helper lineages during adaptive immune responses *in vivo* in settings of chronic inflammation (Gagliani et al., 2015). The suppressive activity of Tr1 cells is predominately due to their production of IL-10 and expression of co-inhibitory receptors (Brockmann et al., 2018; Akdis et al., 2004; Meiler et al., 2008). Indeed, co-expression of one specific co-inhibitory receptor (LAG-3) and an adhesion molecule (CD49b) have been shown to reproducibly identify Tr1 cells in both humans and mice during chronic inflammation and lethal infection (Huang et al., 2017; Brockmann et al., 2018; Gagliani et al., 2013).

Tr1 cells have been shown to inhibit pathogen clearance in models of chronic infection (Heinzel et al., 1991; Ghalib et al., 1993; Oca et al., 2016) and these cells also dampen immune responses during allograft survival and in autoimmunity (Batten et al., 2008; Gagliani et al., 2013; Yu et al., 2017; Karwacz et al., 2017; Zhang et al., 2017). Much less is known about the contribution of Tr1 cells in resolution of acute infection as the infectious models used to date to study these cells have involved lethal infection thereby preventing an analysis of Tr1 cells during response resolution (Huang et al., 2017, 2018; Liu et al., 2014). Thus, it is currently unknown whether Tr1 cells develop as a normal part of T cell responses during acute resolving infection and whether they contribute to immune regulation in this setting. To begin to address these questions, we investigated Tr1 cells in a mouse model of acute resolving respiratory infection with IAV.

## Methods

### Mice

C57BL/6 FOXP3^RFP^IL-10^GFP^ (C57BL/6-*Foxp3*^tm1Flv^ x B6.129S6-*Il10*^tm1Flv^) (Zhang et al., 2017) bred at the University of Adelaide animal house. C57BL/6 *Il27ra*^*-/-*^ (B6N.129P2-*Il27ra*^tm1Mak^ mice) and genetically-matched WT control mice were inter-crossed at the University of Adelaide animal house for littermate experimental mice. C57BL/6J and B6.SJL *Ptprca* (Ly5.1) mice were purchased from the ARC (Perth, WA). All mice were housed under specific pathogen-free conditions at University of Adelaide Animal Facility. Experiments used gender- and age-matched mice between 8 and 12 weeks of age. Mice were humanely euthanised by CO_2_ asphyxiation. All experimental and breeding protocols were approved by the University of Adelaide Animal Ethics Committee under S-2017-040, S-2018-004, S-2019-058 and S-2021-097 ethics approvals.

### IAV infection

X-31 and X-31-OVA_323-339_ [H3N2] IAV stocks were were grown in 10-day-old embryonated chicken eggs. Each egg was injected with 0.1 ml normal saline containing 1 haemagglutination unit (HAU) virus, incubated for 48 h at 37 °C and held at 4 °C overnight. The amniotic/allantoic fluids were then harvested, pooled, clarified and stored at -80 °C. These live virus stocks were titrated in MDCK cells using a TCID50 assay, and the titres were 2.87x10^6^ TCID_50_/mL for X-31 and 3.13x10^5^ TCID_50_/ml for X-31-OVA_323-339_. For IAV infection, mice were anesthetised with pentobarbitone (Ilium) intra-peritoneally (i.p.) and then intranasally (i.n.) instilled with 923TCID_50_ (X-31 IAV) or 200TCID_50_ (X-31-OVA_323-339_) in 32 µL of PBS. Mice were monitored daily for weight loss. Any mouse reaching >20% of initial weight was humanely euthanised as per institutional animal welfare regulations.

### Viral load and qPCR

For viral load analysis whole lungs were snap frozen in liquid nitrogen, ground into fine powder and resuspended in TRIzol (Ambion). RNA was then extracted following the manufacturer’s protocol. RNA samples were DNase treated using Turbo DNA-free kit (Invitrogen), prior to cDNA synthesis (High-Capacity cDNA synthesis Kit, Applied Biosystems). RT-qPCR was conducted using LightCycler-480 SYBR Green I Power-Up Master Mix (Applied Biosystems) on a LightCycler-480 instrument (Roche). Relative expression was calculated by normalising to mouse *Rplp0* as previously described using the 2^-ΔCp^ method (Schmittgen and Livak, 2008). All PCR primers used are listed in **Supplementary Table 1**.

### Flow cytometry, cell sorting, and antibodies

Single cell suspensions were stained in 96 well round-bottom plates (Corning) at 2x10^6^ lymphocytes per well with antibodies and other reagents detailed in **Supplementary Table 2**. Cells were washed once in PBS then resuspended in fixable viability stain 780 (BD) diluted 1/1000 in PBS and blocked with mouse γ-globulin (Rockland) at 10 µg / ml for 5 min at room temperature in the dark. Cells were stained with directly conjugated and biotinylated antibodies diluted in FACS buffer (FB) or Brilliant Stain Buffer (Becton Dickinson) for 20 min at 4°C, with the exception of anti-mouse CD49b (HMα2, BD) and anti-mouse LAG-3 (C9B7W, Biolegend), which were stained for 60 min at 37°C as previously described (Gagliani et al., 2013; Brockmann et al., 2018). For co-staining FOXP3 and IL-10 in WT mice, 2x10^6^ cells/well were restimulated with 10 µg / mL plate bound αCD3 (145-2C11, BioXcell) and 1 µg / mL soluble αCD28 (37.51, BD) and incubated at 37°C and 5% CO_2_. After 2hrs, 20 µL of a solution containing 10 µg /mL of Brefeldin A (eBioscience) and 10 µM Monensin (BD) in complete IMDM were added and mixed gently before incubation for a further 4 hrs at 37°C and 5% CO_2_. After restimulation, cells were transferred to a new plate for staining. Stained cells were analysed using a BD Fortessa X20 cytometer at the University of Adelaide or sorted using a BD FACSAriaIIImu or a BD FACSAria Fusion at The University of Adelaide or SAHMRI.

### *In vitro* suppression assays

5x10^3^ CD11c-enriched APCs were used in each well of the assay, obtained from naïve C57BL/6 spleens using a CD11c positive isolation kit (StemCell). Soluble αCD3 (145-2C11, BioXcell, 1 µg / mL) was added on top. Responding T cells were isolated from spleens and inguinal lymph nodes of naïve Ly5.1 mice using a pan naïve CD3^+^ negative selection kit (StemCell) and then FACS sorted (viable, CD3^+^ CD25^-^) to achieve >95% purity and labelled with Efluor670 proliferation dye (Invitrogen). Tr1 cells were FACS sorted (viable CD3^+^CD4^+^CD44^+^FOXP3^-^IL-10^+^, separated into 4 populations based on co-expression of LAG-3 and CD49b) from IAV-infected FOXP3^RFP^IL-10^GFP^ mouse lungs on day 7 post-infection. 2x10^5^ Tr1 cells were co-cultured with 2x10^5^ labelled T cells for a 1:1 effector: Tr1 ratio. Suppression assays were analysed using the proliferation modelling function in FlowJo (Treestar) and % suppression calculated using the following formula: 100*(1-DI_Tr1_/DI_eff_) based on division index (DI) as described previously (Collison and Vignali, 2011; Ward et al., 2014).

### Intravascular labelling

Three µg of αTCR-β-BV421 (H57-597, Biolegend) was diluted in 200 µL of dPBS (Sigma) per mouse, which was injected via the tail vein 3 minutes prior to humane euthanasia as previously descibed (Anderson et al., 2014).

### Adoptive transfer of Tr1 cells into IAV infection-matched hosts

FOXP3^RFP^IL-10^GFP^ and C57BL/6 mice were infected with X-31 IAV intranasally. On day 7 post-infection, single cell suspensions were prepared from infected lungs from FOXP3^RFP^IL-10^GFP^ mice and cells labelled with efluor670 dye (Invitrogen). Tr1 cells were then FACS-sorted (live CD3^+^CD4^+^CD44^+^FOXP3^RFP-^IL-10^GFP+^ separated into 4 populations based on co-expression of LAG-3 and CD49b). Infection-matched C57BL/6 mice received Tr1 cells or PBS intravenously (i.v.). On day 9 post-infection, recipients were humanely euthanised and tissues harvested to quantify transferred Tr1 cells and assess plasticity.

### RNA sequencing

For sequencing, lung cells were isolated and stained and Tr1 cell populations were FACS-sorted using a FACSAriaIIImu. Total RNA was extracted using a Pico Pure RNA Extraction Kit (Articus) following the manufacturer’s protocol. DNA digestion was performed using an RNase-free DNase Kit (Qiagen). Total RNA was eluted in 20µL elution buffer. RNA sequencing of 150bp paired end reads at an average of 30M reads per sample was performed by Novogene. Data QC for the RNA sequencing experiment was conducted using FastQC (Andrews, 2019) and ngsReports (Ward et al., 2020). Read trimming and filtering was performed using AdapterRemoval v2.2.1 (Schubert et al., 2016), with reads aligned to GRCm38 using STARv2.5.3 (Dobin et al., 2013). Reads were summarised to gene-level counts using featureCounts and gene annotations provided by Ensembl Release 96. Differential Expression analysis was performed using voom (Law et al., 2014) with sample-level quality weights assigned by nesting the four cell lines within each mouse. An FDR-adjusted p-value < 0.05 along with an estimated log Fold-Change (LFC) beyond the range ±1 was used to consider a gene as differentially expressed, Enrichment testing was performed using ClusterProfiler (Yu et al., 2012), taking all detectable genes as the universe against which to test for enrichment. Mapping from gene to Gene Ontology was obtained using the Bioconductor package org.Mm.eg.db, and a Bonferroni-adjusted p-value < 0.05 was used to consider an ontology enriched in any of the sets of DE genes, (Kolde, 2012). An UpSet plot was generated from a complete list of DE genes after filtering by FDR and LFC. For this comparison any genes with an FDR-adjusted p value <0.05 was considered significant in this analysis to give a comprehensive result and define population-specific signature genes. All analytic code is available via github (https://caitlinabbott.github.io/Tr1-RNA-Sequencing-/).

## Results

### Tr1 cells emerge during acute viral infection and contribute to resolution of the response

The significance of Tr1 cells in resolution of acute viral infection has not been explored to date. To address this, we utilised intranasal installation of X-31 H3N2 IAV. In C57Bl/6 mice, this infection leads to transient viral replication in the lungs associated with moderate weight loss that completely resolves by day 10 post-infection (**Fig. S1**). T cells that secrete IL-10 in the lung in this setting were identified and assessed by flow cytometry using FOXP3-RFP mice (Wan and Flavell, 2005) crossed to IL-10-GFP (TIGER) (Kamanaka et al., 2006) mice, referred to hereafter as FOXP3^RFP^IL-10^GFP^ mice (**Fig 1A**). Prior to infection, and at day 5 post-infection, the majority of T cell-derived IL-10 in lungs emanated from innate-like (CD4^-^CD8^-^) cellular sources (**Fig 1B, C**). However, by day 7, CD4^+^ T cells were clearly the major T cell subset responsible for IL-10 production, outnumbering IL-10^+^ CD8^+^ T cells approximately 2:1 (**Fig 1C**) and producing significantly more IL-10 than other T cells in the lung on a per cell basis by mean fluorescence intensity (MFI) (**Fig 1D**). Although there was some contribution from Tregs, defined here as CD4^+^ FOXP3^+^, to IL-10 production, the predominant CD4^+^ T cell subset in the lung producing IL-10 was FOXP3^-^, consistent with Tr1 cells (**Fig 1E-G**). These cells did not express *Il4, Il17a, Il22, or Csf1* but did express *Ifng, Tgfb*, and *Il21* (**Fig. S2**), features that are in line with a Tr1 cytokine signature (Bacchetta et al., 1990; Gagliani et al., 2013).

**Figure 1:**
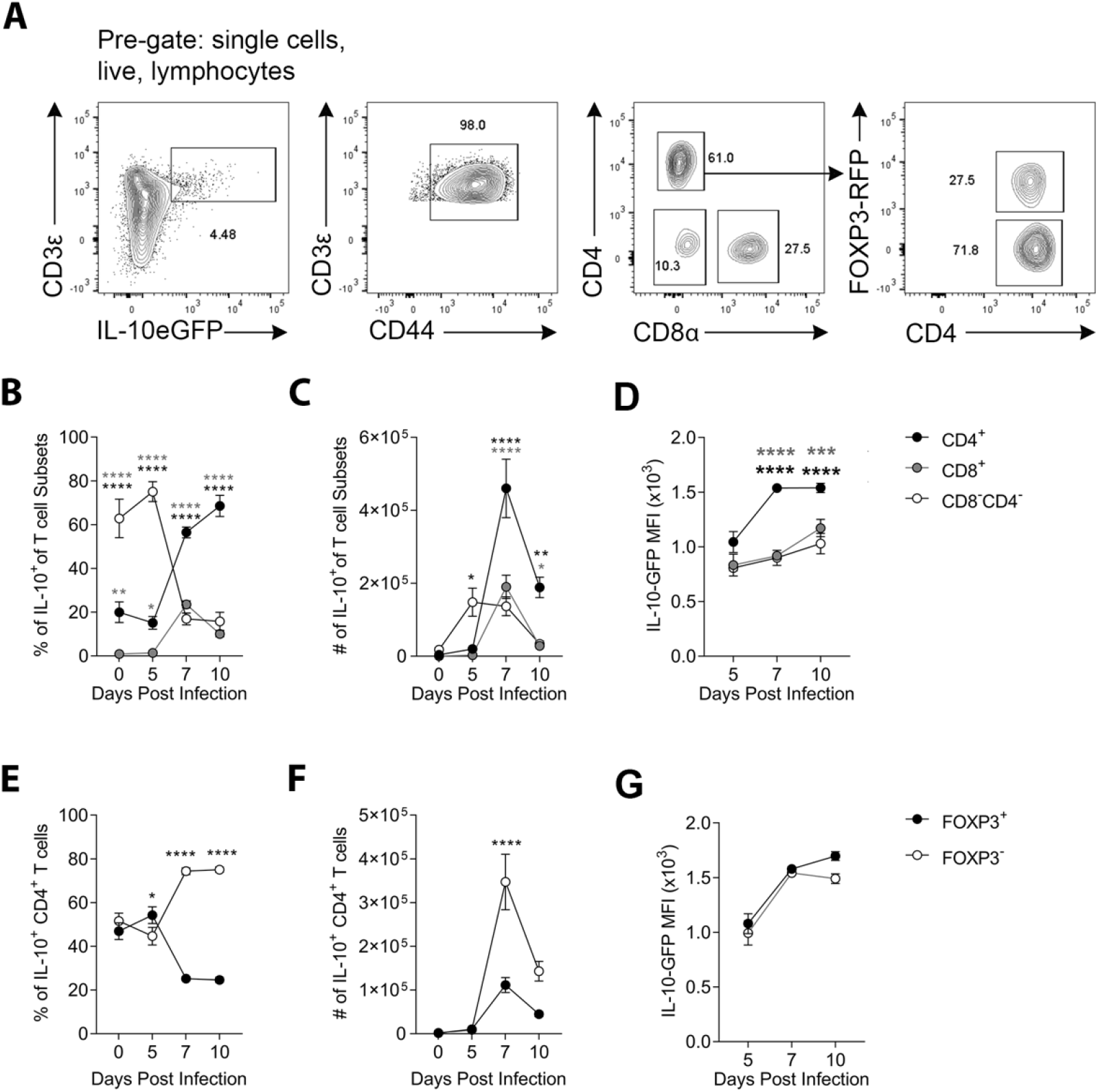
Tr1 cells are the dominant source of T cell-derived IL-10 during IAV-infection. Dual-reporter mice were infected with X-31 IAV-i.n. Activated T cell populations from the lungs were analysed at days 0, 5, 7, and 10 post-infection. (A) Representative gating for identification of IL-10^+^ CD3^+^ T cell populations from IAV-infected lungs. (B) Frequency, (C) number, and (D) MFI of IL-10-secreting CD8^+^, CD4^+^, and CD8^-^ CD4^-^ T cells. (E) Frequency, (E) number, and (F) MFI of IL-10-producing FOXP3^-^ (Tr1 cells) and FOXP3^+^ (Treg cells) over the course of IAV infection. (B, C, E, F) The mean is shown +/- SEM, n=7 (d0), 9 (d5), 13 (d7), 10 (d10) biological replicates total from 3 independent experiments. Statistical analysis using one-way ANOVA with Bonferroni’s post-test. (D) The mean is shown +/- SEM, n=6 (d5), 7 (d7), 7 (d10) biological replicates total from 6 independent experiments. Statistical analysis using 2-way ANOVA with Bonferroni’s post-test * p ≤ 0.05, ** p ≤ 0.01, *** p ≤ 0.001, **** p ≤ 0.0001.

To establish the kinetics of CD4^+^ T cell accumulation during IAV infection, the abundance of activated effector cells (CD44^+^FOXP3^-^IL-10^−^), Tregs (CD44^+^FOXP3^+^IL-10^−^), IL-10^+^ Tregs (CD44^+^FOXP3^+^IL-10^+^), and Tr1 cells (CD44^+^FOXP3^-^IL-10^+^) were quantified during the course of infection in the spleen, lung-draining mediastinal lymph node (mLN), lungs, and peripheral blood (**Fig 2**). As expected, activated CD4^+^ effector T cells were the most abundant of these populations in each of the different compartments (**Fig 2A-E**). In order to interrogate the differences in accumulation between the different regulatory T cell populations (Tregs, IL-10^+^ Tregs, and Tr1 cells) these populations were directly compared (**Fig 2A-E**). This revealed that Tregs increased in the mLN and spleen between day 5 and 7 post-infection, however, this was not observed for Tr1 cells or IL-10^+^ Tregs (**Fig 2A, B-C**). However, both Tregs and Tr1 cells had accumulated to a similar extent by day 7 in the lungs (**Fig 2A, D**). In addition, compared to Tregs, neither Tr1 cells nor IL-10^+^ Tregs substantially increased in frequency in the blood **(Fig 2A, E)**. Together, these findings are consistent with limited generation of Tr1 cells and IL-10^+^ Tregs in secondary lymphoid organs (SLO) and instead support the notion that both Tr1 cells and IL-10^+^ Tregs are predominately generated from infiltrating precursors at the site of infection.

**Figure 2:**
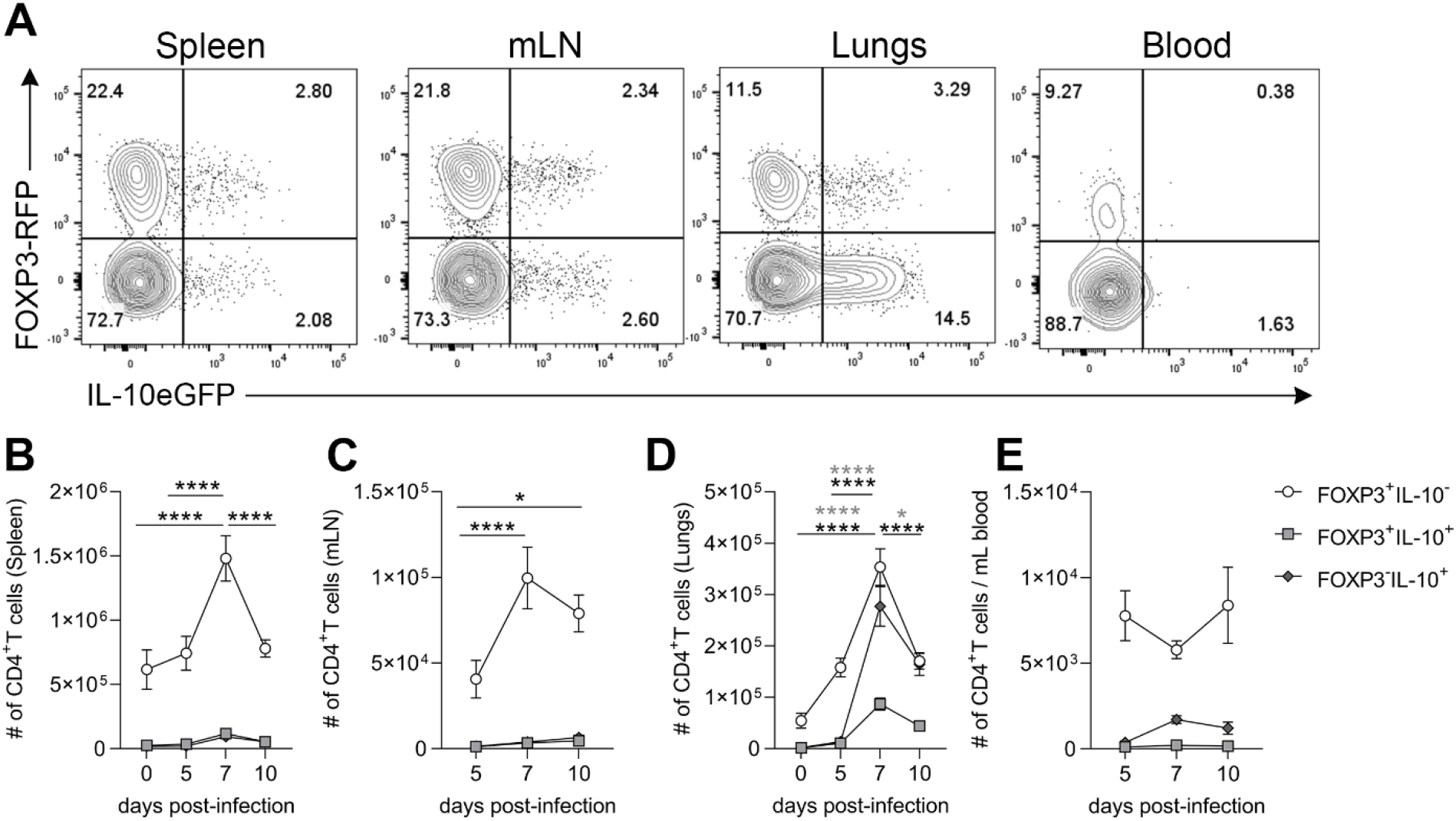
Tr1 cells transiently accumulate in the lung during IAV infection without significant expansion in SLOs. Dual-reporter mice were infected with X-31 IAV-i.n and on day 5, 7, and 10 post-infection spleens, mLN, lungs and peripheral blood were harvested and activated (CD44^+^) CD4^+^ T cell populations were quantified by flow cytometry. (A) Representative flow cytometry of CD4^+^ T cell populations across all four organs on day 7. The number of each CD4^+^ T cell population across the time-course in the (B, F) spleen, (C, G) mLN, (D, H) lungs, (E, I) per mL of peripheral blood. Each symbol represents the mean +/- SEM. For spleens n=7-22, mLNs n=10-12, lungs n=7-29 and PB n=7-10, all pooled from 7 independent experiments. Statistical analysis using 2-way ANOVA with Bonferroni’s post-test where *p<0.05, **p<0.01, ****p<0.0001.

After establishing that Tr1 cells transiently accumulated in the lung at day 7, which coincided with the beginning of response resolution **(Fig S1A)**, the contribution of Tr1 cells to the resolution of acute IAV infection was assessed. *Il27ra*^*-/-*^ mice were utilized, which have been reported to have a selective deficiency in the generation of Tr1 cells but not in FOXP3^+^ Tregs(Batten et al., 2008). *Il27ra*^*-/-*^ mice lost more weight than littermate controls, exhibited a delay in post-infection weight recovery (**Fig 3A**), and had a selective loss of Tr1 cells in the lung (**Fig 3B-G**) on day 7 of the response. There were no deficits in either IL-10^+^ or IL-10^−^ Tregs between *Il27ra*^*-/-*^ and control mice (**Fig 3B-D**). In fact, there was a significant increase in FOXP3^+^ IL-10^−^ Tregs, potentially compensating for observed deficiency of Tr1 cells. In the BALF of WT mice, there was an enrichment of all IL-10^+^ CD4^+^ T cells compared to the lungs, suggesting IL-10^+^ cells preferentially localise in the airways (**Fig 3E-G**). Similar to the lungs, there was no reduction in Tregs between *Il27ra*^*+/+*^ and *Il27ra*^*-/-*^ mice in the BALF. However, there was a stark ablation of Tr1 cells in the BALF of *Il27ra*^*-/-*^ mice on day 7 post-infection (**Fig 3E-G**). Together, these data suggest a role for IL-10-secreting Tr1 cells in the resolution of acute IAV infection of the lung.

**Figure 3:**
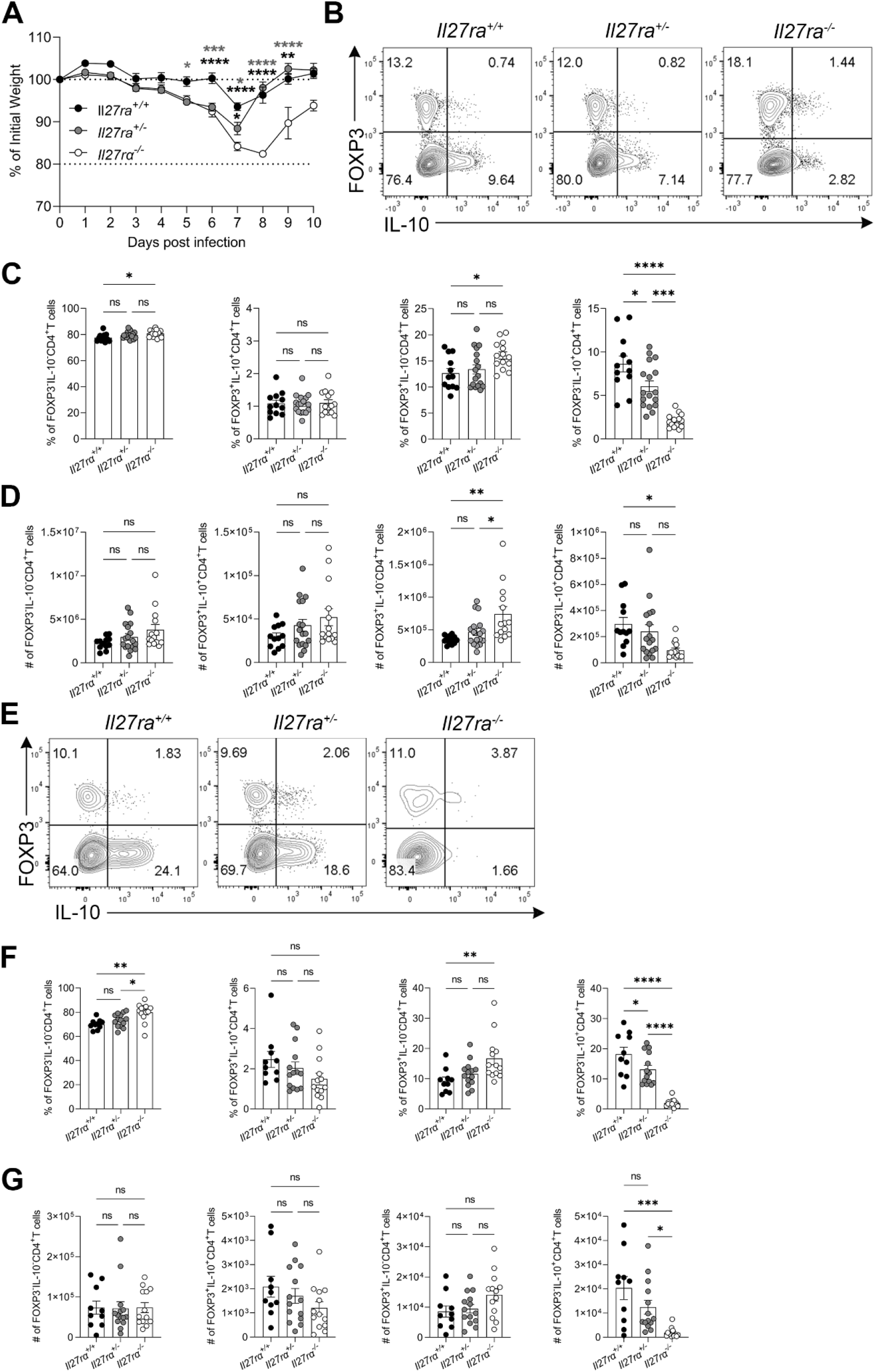
*Il27ra*^*-/-*^ mice exhibit a deficiency in Tr1 cells and have delayed resolution of acute IAV infection. *Il27ra*^*+/+*^, *Il27ra*^*+/-*^, and *Il27ra*^*-/-*^ mice were infected with X-31-OVA_323-339_ IAV i.n and lungs were analysed on day 7 post-infection. (A) The percentage of initial weight over the course of IAV infection. (B) Representative flow cytometry of activated CD4^+^ T cells in the lungs at day 7 post-infection. (C) Percentage and (D) number of CD4^+^ T cell populations in the lungs on day 7 post-infection. (E) Representative flow cytometry of activated CD4^+^ T cells in the BALF at day 7 post-infection. (F) Percentage and (G) number of CD4^+^ T cell populations in the BALF on day 7 post-infection. (A) The mean is shown +/- SEM. (C-D) each symbol represents a different biological replicate, shown as mean +/- SEM, n=12 (*Il27ra*^*+/+*^), 17 (*Il27ra*^*+/-*^), 14 (*Il27ra*^*-/-*^) biological replicates total from 6 independent experiments. Statistical analysis using (A) 2-way ANOVA, (C, D, F, & G) one-way ANOVA both with Bonferroni’s correction * p ≤ 0.05, **p ≤ 0.01, ***p ≤ 0.001, ****p ≤ 0.0001.

### Tr1 cells in IAV infection have a heterogeneous LAG-3 and CD49b surface phenotype and are localised in the parenchyma of the lungs

To explore the biology of Tr1 cells in the context of acute IAV infection in more detail, we examined the expression of LAG-3 and CD49b (**Fig. S3**), which have previously been described as characteristic surface markers of Tr1 cells (Brockmann et al., 2018; Huang et al., 2017; Gagliani et al., 2013). In contrast to Tr1 cells described in chronic inflammatory settings (Brockmann et al., 2018; Gagliani et al., 2013), Tr1 cells in acute IAV infection were predominately double negative (DN, LAG-3^-^CD49b^-^) during the early phase of the response (day 5 post-infection). By day 7, four populations of Tr1 cells were distinctly apparent based on LAG-3 and CD49b co-expression **(Fig. 4A)**. These included Tr1 cells that resemble canonical Tr1 cells (double positive (DP) LAG-3^+^CD49b^+^), but also substantial populations that were single positive for either LAG-3^+^ or CD49b^+^. Tr1 cells that expressed at least one of these markers increased in the lung until day 7 post-infection, after which they markedly dropped in frequency (**Fig. 4B**). In contrast, DN Tr1 cells continued to accumulate in lung up to day 10 post-infection (**Fig. 4B**). Tr1 cell co-expression of IL-10 with IFNγ has been shown to align with LAG-3 and CD49b co-expression in some studies (Levings et al., 2001; Brockmann et al., 2017). Therefore, we also interrogated activated FOXP3^-^ CD4^+^ T cells in IAV, subdivided on the basis of CD49b and LAG-3 co-expression and found that all 4 populations contained a population of IL-10^+^IFNγ^+^ cells, but that LAG-3 single-positive and DP populations were similarly enriched for co-expression of these cytokines (**Fig. S4**). Together, these data define that a heterogeneous population of Tr1 cells emerges during acute IAV infection. These include LAG-3^+^ CD49b^+^ (DP) cells consistent with the previously described Tr1 cell phenotype, although a majority display an atypical phenotype where LAG-3 and CD49b are not co-expressed.

**Figure 4:**
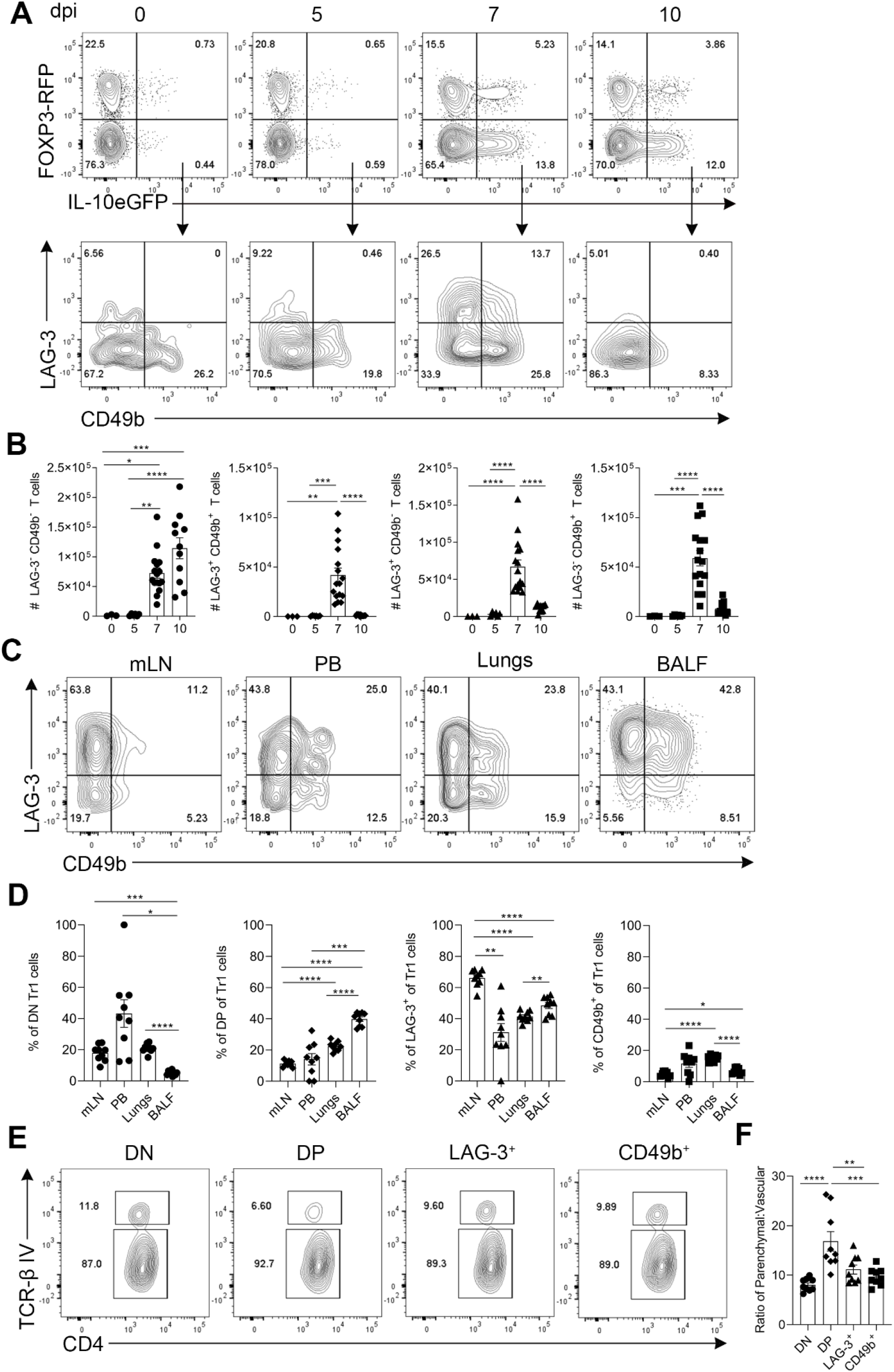
IAV-induced Tr1 cells exhibit heterogeneous expression of LAG-3 and CD49b and are localised in the lung parenchyma. Dual-reporter mice were infected with X-31 IAV i.n. (A) On days 0, 5, 7, and 10 post-infection mice were humanely killed, and lungs were harvested. Concatenated representative flow cytometry depicting expression of FOXP3 and IL-10 by activated CD4^+^ T cells and LAG-3 and CD49b by Tr1 cells. (B) The number of LAG-3^+^ CD49b^-^, LAG-3^+^ CD49b^+^, LAG-3^+^ CD49b^-^, and LAG-3^-^ CD49b^+^ Tr1 cells post IAV infection. (C) Concatenated representative flow cytometry from the indicated organs at day 7 post infection, displaying LAG-3 and CD49b expression by Tr1 cells. (D) Frequency of DN, DP, LAG-3^+^ and CD49b^+^ Tr1 cells in the mLN, PB, lungs, and BALF. (E) Concatenated flow cytometry depicting TCR-β I.V labelling (top TCR-β^+^ gate in E) of Tr1 cells from the lungs on day 7 post-infection. (F) The ratio of Tr1 cells in the parenchyma vs the vasculature. Each symbol represents a different biological replicate, bars show the mean, and error bars depict +/- SEM, (A-B) n=3-15 biological replicates total per time point from 3 independent experiments, (C-F) n=9 biological replicates total from 2 independent experiments. Statistical analysis using (B, F) one-way ANOVA with Bonferroni’s post-test, and (D) paired Student’s T-tests * p ≤ 0.05, ** p ≤ 0.01, *** p ≤ 0.001, **** p ≤ 0.0001.

We next investigated the tissue distribution of these 4 Tr1 cell phenotypes following acute IAV infection. The peripheral blood, mLN, lungs, and BALF were examined for the presence of Tr1 cells separated on the basis of LAG-3 and CD49b on day 7 post-infection. Strikingly, LAG-3 single-positive cells were the most abundant Tr1 population in each of these compartments (**Fig. 4C, D**). Interestingly, DP Tr1 cells were more frequent in the BALF than in other tissues examined suggesting these cells may be preferentially expanded, recruited, or maintained within the airways of infected lungs. Next, we performed intravascular labelling to determine whether these Tr1 cell phenotypes differentially occupied the parenchyma or the vasculature of the lungs. Greater than 80% of Tr1 cells were found within the parenchyma of the IAV-infected lungs on day 7 post-infection (**Fig. 4E**). Each of the four Tr1 phenotypes were enriched in the parenchyma compared to the vasculature of the lungs (**Fig. 4E**). However, the DP population was the most enriched in the parenchymal niche amongst the Tr1 cells in IAV infection (**Fig. 4F**), suggestive of a later stage of tissue infiltration.

### Tr1 cells from IAV-infected lungs are *bona fide* Tr1 cells

The heterogeneity observed with respect to LAG-3 and CD49b expression in the Tr1 compartment in acute IAV infection raised the question of whether all of these apparent Tr1 cell phenotypes are *bona fide* suppressor cells, capable of inhibiting T cell division. To address this, we performed suppression assays using labelled effector T cells. Tr1 cells of each of the 4 identified phenotypes were sorted from lungs of IAV-infected FOXP3^RFP^IL-10^GFP^ mice on day 7 post-infection for these assays **(Fig. S5)**. All four Tr1 cell populations were capable of significant titratable inhibition of effector T cell division **(Fig. 5A-C)**, with the DP and LAG-3^+^ populations most suppressive on a per cell basis (significantly suppressive out to a 1:8 Tr1:effector T cell ratio) **(Fig. 5C)**. These results indicate that all 4 Tr1 cell phenotypes in IAV are suppressive and can be considered *bona fide* Tr1 cells with the DP and LAG-3^+^ populations perhaps exhibiting the most potent regulatory activity. Mechanistically, Tr1 cells have been reported to exert suppression predominately via IL-10 production (Groux et al., 1997; Gagliani et al., 2013; Gregori et al., 2010). Thus, each of the four Tr1 cell populations were tested for their dependence on IL-10 to elicit suppression by conducting assays with a neutralising α-IL-10Rα antibody **(Fig. 5D, E)**. DP and CD49b^+^ Tr1 cells exhibited some dependence on IL-10 to elicit suppression, while DN and LAG-3^+^ Tr1 cells suppressed effector T cell division independently of IL-10 **(Fig. 5D, E)**. Although IL-10 was important for suppression exerted by DP and CD49b^+^ populations of Tr1 cells, this indicated that other mechanisms were employed by LAG-3^+^ and DN Tr1 cells to inhibit effector T cell division.

**Figure 5:**
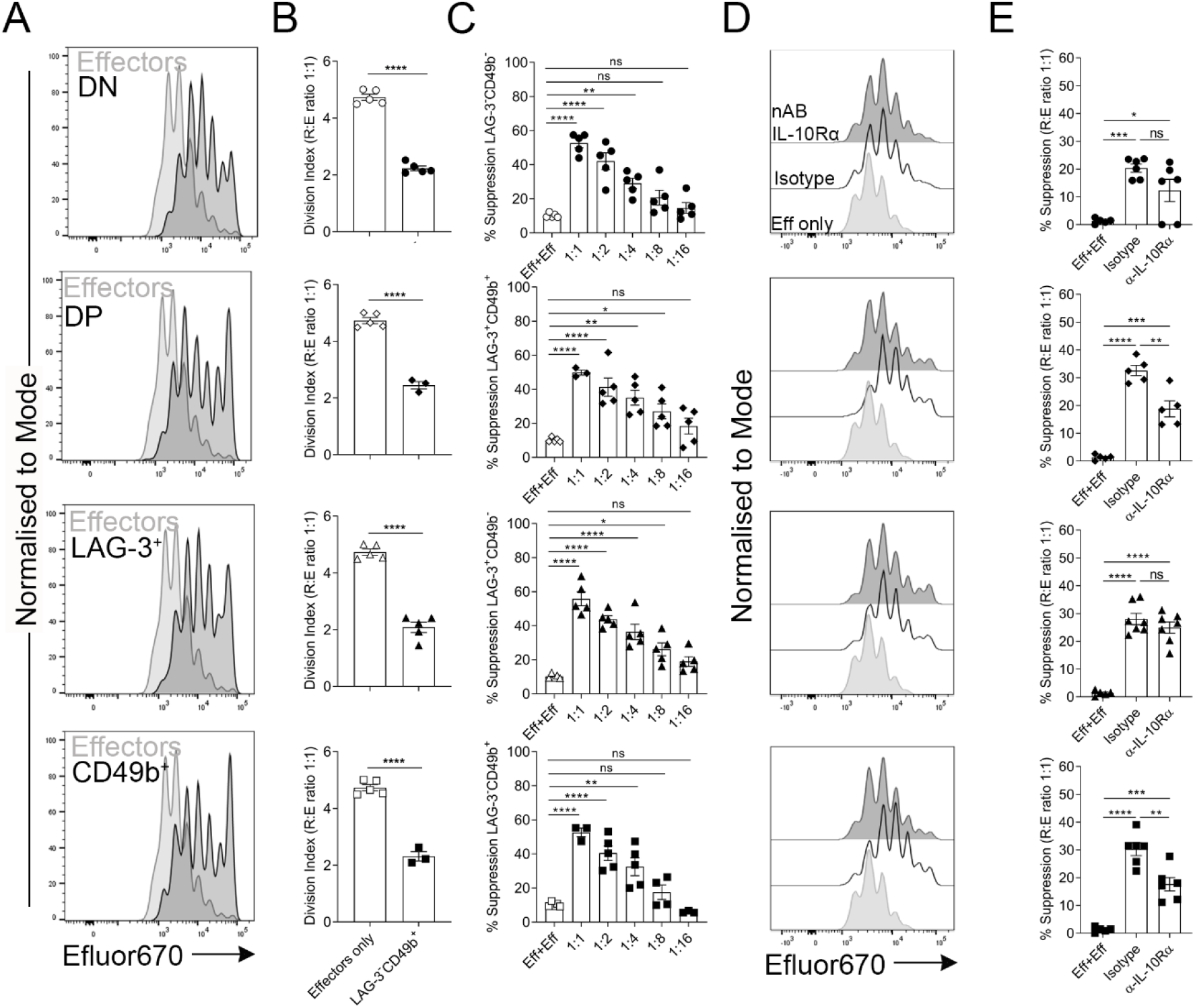
FOXP3^-^ IL-10^+^ CD4^+^ T cells induced by IAV infection are *bona fide* Tr1 cells capable of suppressing effector T cell division. Dual-reporter mice were infected with X-31 IAV i.n. and Tr1 cell populations were FACS-sorted from lungs on day 7 post-infection for suppression assays. (A) Representative flow cytometry showing the proliferation of Efluor670-labelled CD3^+^ CD25^-^ responder T cells with stimulation alone (light grey histograms, Effectors) or in the presence of Tr1 cells (overlaid dark grey histogram) at a 1:1 ratio (2x10^4^ Tr1 cells: 2x10^4^ effector T cells). (B) Division index (DI) of effector T cells cultured with APCs and α-CD3 alone or with Tr1 cells at a 1:1 ratio (where DI= the mean number of divisions within a population). (C) The percentage suppression of effector T cell division at each ratio of Tr1 cells: effector T cells (1:1-1:16) compared to effector T cells cultured with effector T cells (Eff+Eff). (D) Representative flow cytometry depicting division of labelled effector T cells from suppression assays. Stimulation alone condition (light grey histograms), co-cultured with each of the Tr1 cell populations (1:1 ratio) in the presence of an isotype control (black histograms), in the presence of IL-10Ra nAB (dark grey histograms). Data are shown as mean +/- SEM, (A)-(C) n=3-5 biological replicates, (D) & (E) n=4-7 replicates from 3 independent experiments. Statistical analysis using (B) an unpaired Student’s T-test and (C, D) one-way ANOVA with Bonferroni’s post-test * p < 0.05, ** p < 0.01, *** p < 0.001, ****p < 0.0001.

### Transcriptomic analysis of lung derived Tr1 populations

Given the differences observed in kinetics, localisation, and dependence on IL-10 for suppressive capacity between the 4 Tr1 cell populations identified in IAV infection, further differences were probed at the transcriptional level. RNA sequencing was conducted on each of the four populations sorted from lungs of IAV-infected FOXP3^RFP^IL-10^GFP^ mice on day 7 post-infection **(Fig. S6)**. Using an FDR <0.05 and estimated log-fold-change (logFC) greater/less than 1(+/-1), 243 differentially expressed (DE) genes were identified in total (where each unique gene was determined to be DE in at least one comparison). An UpSet plot was generated from a list of the DE genes across all 6 comparisons. This analysis revealed that the LAG-3^+^ and CD49b^+^ populations exhibited 169 DE genes, the greatest number in any one comparison and consequently appeared the most distinct among all the populations **(Fig 6A)**. Unique gene set signatures were determined for the DN, LAG-3^+^, and CD49b^+^ Tr1 cell populations but not for the DP Tr1 cells **(Fig 6A, Fig S7A)**. This is likely due to the DP sharing features of both the LAG-3^+^ and CD49b^+^ Tr1 cells meaning there are no significant uniquely expressed genes by this population. Next, it was established how closely each of the four Tr1 cell populations from acutely infected mice resembled previously reported Tr1 cells and other CD4^+^ T cell subsets. To address this, a list of Tr1 cell signature genes was compiled from previous studies (Pot et al., 2009; Brockmann et al., 2018; Gagliani et al., 2013; Karwacz et al., 2017; Zhang et al., 2020). We compared the expression of Tr1 signature genes as well as genes associated with Th1, Th2, Th17, Tfh and Tregs between the four Tr1 cell populations from IAV-infected lungs. Apart from the genes encoding the surface markers used for their differential sorting, it was apparent that all four populations of Tr1 cells strongly conform to a core Tr1 cell gene expression signature (i.e. abundant *Il10, Maf, Gzmb, Icos, Tigit, Ctla4*, and *Pdcd1*) **(Fig. 6B)**. An exception amongst these Tr1 cell-associated genes was *Eomes*, which encodes a transcription factor that has been shown to control Tr1 cell development during chronic alloantigen exposure in GVHD (Zhang et al., 2017). In acute IAV, *Eomes* was not highly expressed in Tr1 cells, as would be expected in an acute setting, but was present at low levels only in the two CD49b^-^ Tr1 cell populations. This was confirmed by staining intracellular EOMES protein in these cells (**Fig. S8**). In addition, acute IAV infection-induced Tr1 cells generally did not express high levels of genes associated with Th2 (except *Gata3*), Th17, Tfh or Tregs (except *Il2ra* and *Il2rb*) but did abundantly express Th1-associated genes including *Tbx21, Cxcr3, Cxcr6* and *Ifng*. Together, these data indicate that the atypical Tr1 cell phenotypes that arise during acute IAV infection strongly resemble previously described Tr1 cells and that the LAG-3^+^ and CD49b^+^ phenotypes were the most transcriptionally distinct amongst these populations.

**Figure 6:**
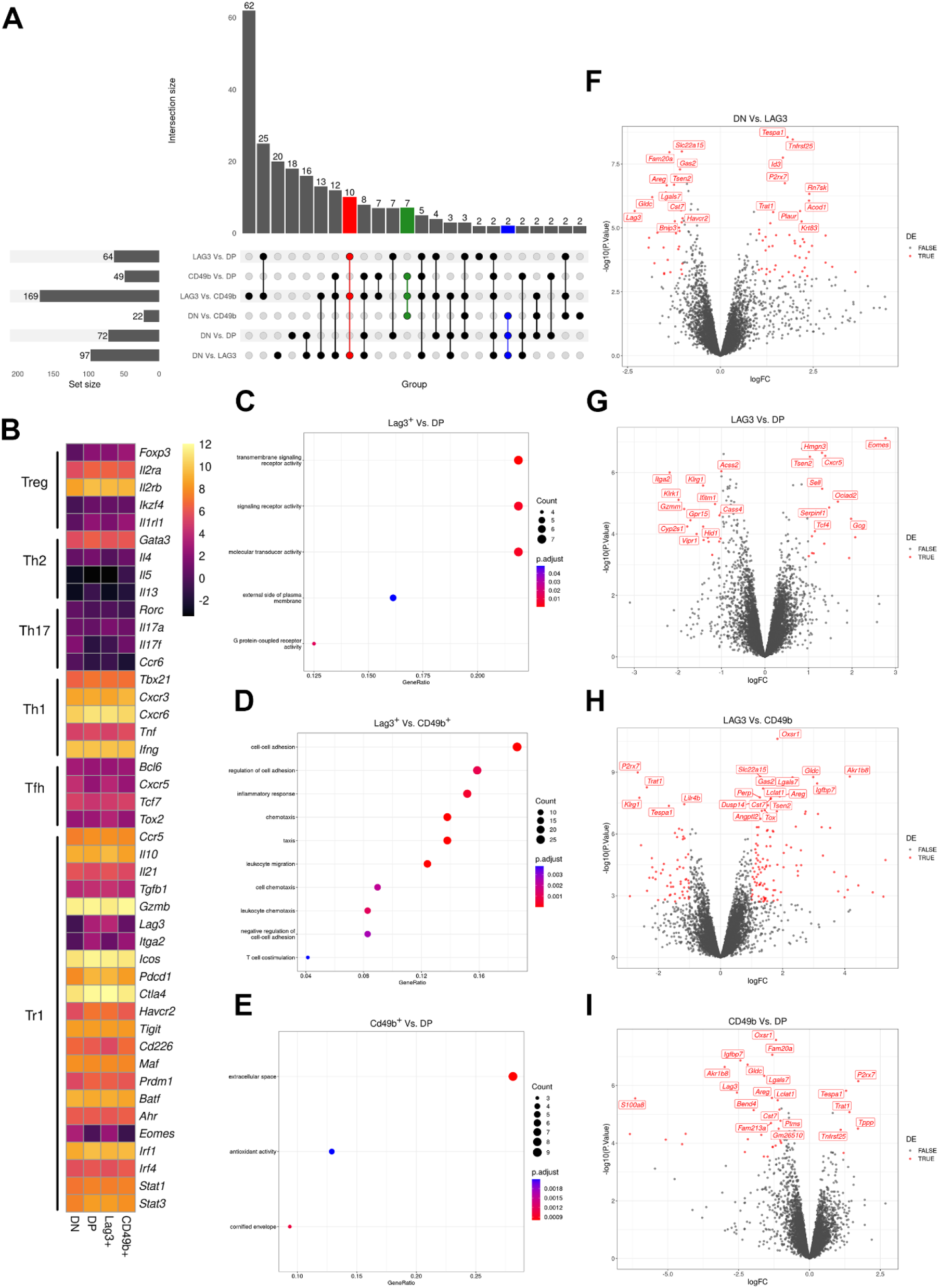
Four populations of IAV-induced Tr1 cells conform to a general Tr1 cell gene expression signature but exhibit subset-specific transcriptional divergence. Dual-reporter mice were infected with X-31 IAV i.n. and on day 7 post-infection DN, DP, CD49b+ and LAG-3+ subsets of Tr1 cells were FACS-sorted and analysed by RNA-sequencing. (A) UpSet plot for all DE genes. (B) Heatmap depicting average mRNA expression of Tr1, Treg, Th2, Th17, Th1, and Tfh lineage signature genes across the four Tr1 populations (generated using pheatmap in RStudio). The heatmap scale refers to the average logCPM on a continuous scale for each gene listed. (C-F) Pathway analysis was conducted using clusterprofileR in R studio and dot plots are shown for the comparisons between (C) LAG-3^+^ vs DP, (D) LAG-3^+^ vs CD49b^+^ and (E) CD49b^+^ vs DP. Dot size is proportional to the number of genes in a given EnterezID gene set and a Bonferroni-adjusted p <0.05 was used as a cut off for significantly enriched pathways. (F-I) Volcano plots were generated using ggplot2 in R studio with comparisons between (F) DN vs Lag3^+^, (G) Lag3^+^ vs DP (H) Lag3^+^ vs CD49b^+^ and (I) CD49b^+^ vs DP. The top 20 differentially expressed genes are labelled in each comparison. Coloured dots indicate significantly differentially expressed genes based on a false discovery rate (FDR) < 0.05 and an estimated logFC greater than/less than +/- 1. N=4 biological replicates for each population from two independent experiments were analysed for RNA sequencing.

Next, to further explore differences between Tr1 cell phenotypes, pathway analysis was conducted to identify the gene ontologies (GO) significantly enriched in each comparison. First, DN and LAG-3^+^ Tr1 cells were compared, and this revealed 58 GO terms significantly enriched including ‘inflammatory response’ highlighting genes such as *Havcr2, Cst7*, and *Lag3* expressed by the LAG-3^+^ Tr1 cells **(Fig S7C)**. Between the LAG-3^+^ and DP Tr1 cells, 3 significantly enriched GO terms were identified including ‘transmembrane receptor activity’ (e.g. *Cxcr5, Itga2, Klrk1, and Gpr15*) **(Fig 6C)**. DP Tr1 cells expressed more *Itga2, Klrk1*, and *Gpr15*, whereas LAG-3^+^ Tr1 cells expressed more *Cxcr5*. This suggested the DP Tr1 cells expressed more genes associated with T cell activation and adhesion than LAG-3^+^ Tr1 cells and therefore potentially constitute a more differentiated population. The comparison between LAG-3^+^ and CD49b^+^ Tr1 cells identified 47 significantly enriched GO terms including ‘cell-cell adhesion’ (e.g. *Lgals7, Perp, Tespa1, Lag3, Itgb3, Itga2*, and *Il10*) **(Fig 6D)**. This analysis indicated that CD49b^+^ Tr1 cells expressed increased levels of integrins and exhibited a more terminally differentiated gene signature than LAG-3^+^ Tr1 cells. Between CD49b^+^ and DP Tr1 cells only 3 GO terms were considered significantly enriched. These included ‘extracellular space’ (e.g. *Fam20a, Igfbp7, Lgals7, Areg, S100a8, Cst7, Csf1, S100a9*, and *Pla1a*) **(Fig 6E)**. The ‘extracellular space’ GO term encompassed genes encoding surface and secreted molecules that were DE by the DP Tr1 cells and could thus be attributed a more activated and functional cell phenotype. Collectively, the pathway analysis identified differences in GO terms associated with activation and adhesion suggesting that the major differences in these Tr1 populations are related to their activation and functional state with DP cells potentially representing the most activated phenotype.

Upon examining expression of specific genes between the 4 Tr1 populations in more detail it was apparent that LAG-3^+^ Tr1 cells exhibited a more activated and regulatory profile than DN Tr1 cells (e.g. increased expression of *Havcr2, Lag3*, and *Areg*) **(Fig 6F)**. Indeed, compared to each of the other populations DN Tr1 cells appeared more Tfh-like and less regulatory **(Fig. 6F, Fig. S7)**. Comparison between LAG-3^+^ and DP Tr1 cells revealed DE genes upregulated in LAG-3^+^ Tr1 cells that were suggestive of a less differentiated state (e.g. *Sell* and *Cxcr5*) **(Fig 6G)**. In contrast, genes increased in DP Tr1 cells were consistent with a more activated and functional phenotype than the LAG-3^+^ Tr1 cells (e.g. DE genes included *Klrk1, Klrg1, Gzmm*, and *Itga2*) **(Fig 6G)**. Consistent with the observations from the GO term analysis, compared to both LAG-3^+^ and DP cells, CD49b^+^ Tr1 cells exhibited more features of active TCR-signalling (e.g. *Tespa1* and *Trat1*) in combination with increased expression of receptors associated with induction of cell death (e.g. *P2rx7* and *Tnfr25*) **(Fig 6H-I)**, which implied a more apoptotic phenotype of CD49b^+^ Tr1 cells. Thus, there are differences apparent in the expression of genes associated with T cell activation and fitness between the various Tr1 cell populations from lungs of acutely infected mice.

### Tr1 cell populations exhibit differences in activation and fitness

As genes associated with activation and death were highly differentially expressed between subsets of Tr1 cells, several of these were probed further. TIM-3 and PD-1 (activation markers induced upon TCR signalling and activation) (Agata et al., 1996; Hastings et al., 2009) were first selected for further interrogation. Tr1 cells express a range of co-inhibitory receptors (Brockmann et al., 2018), which have been shown to contribute to their suppressive capacity in a number of different contexts (Groux et al., 1997; Vieira et al., 2004; Chen et al., 2021). Expression of such receptors is also an indication of activation and antigen experience thereby allowing their co-expression to provide a readout of the activation level of Tr1 cells. Both DP and LAG-3^+^ Tr1 cells expressed significantly greater surface PD-1 compared to DN and CD49b^+^ Tr1 cells **(Fig 7A, B)**. Furthermore, the DP Tr1 cells expressed the highest level of surface TIM-3 and exhibited the greatest proportion of PD-1^+^ TIM-3^+^ double-positive cells **(Fig 7A, D)**, further supporting the notion that DP Tr1 cells are the most activated of the four populations.

**Figure 7:**
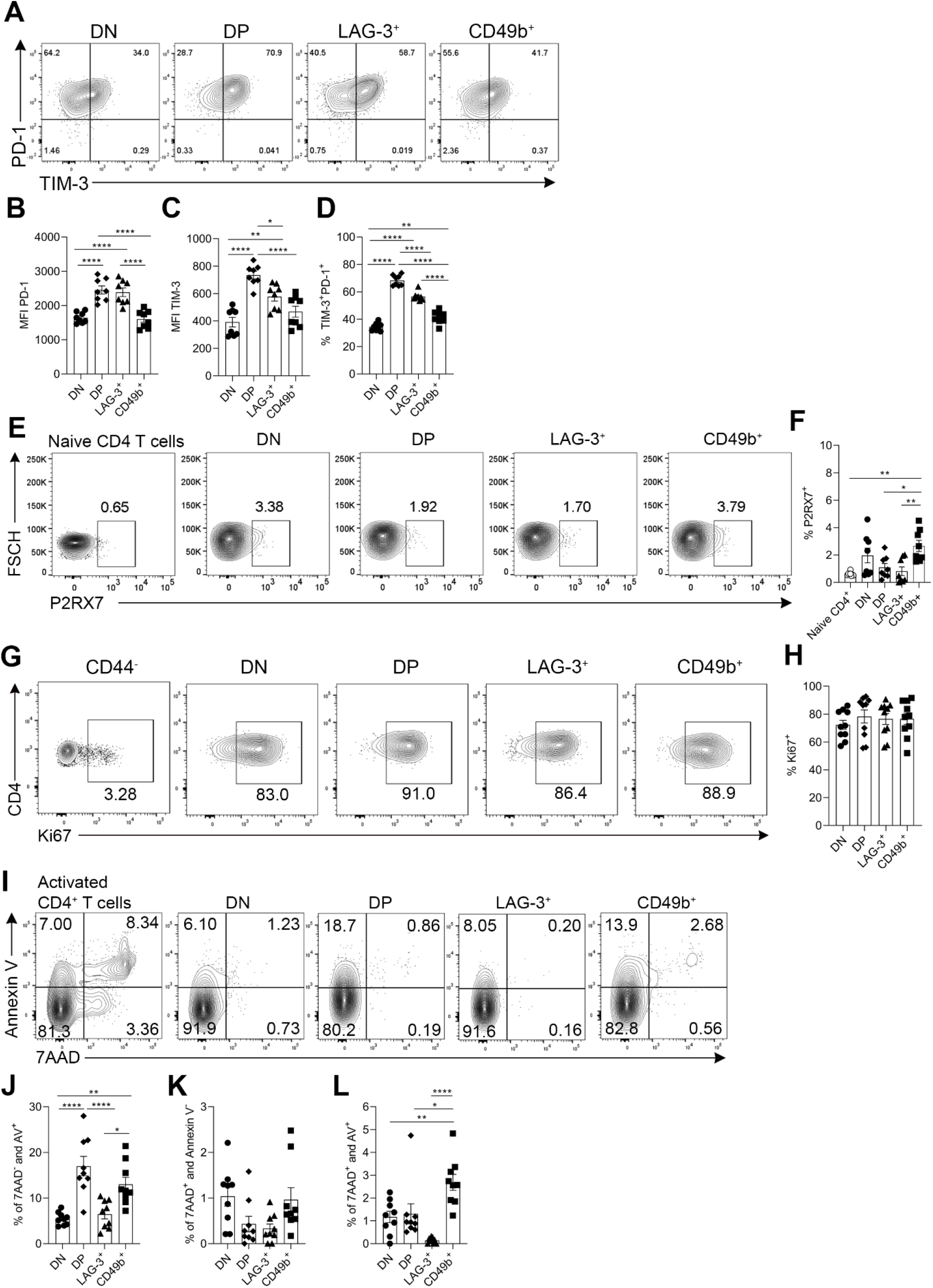
Activation, proliferation, and death of Tr1 cell populations. Dual-reporter mice were infected with X-31 IAV-i.n. Lungs were harvested for on day 7-post infection. (A) Representative flow cytometry of TIM-3 and PD-1 co-expression by Tr1 cells. MFI of (B) PD-1 and (C) TIM-3. (D) The percentage of PD-1 TIM-3 double positive cells. (E) Representative flow cytometry of P2RX7 expression by each of the four Tr1 cell populations compared to naïve CD4^+^ T cells. (F) Frequency of P2RX7^+^ cells. (G) Representative flow cytometry depicting expression of intra-nuclear Ki67 by Tr1 cells. (H) Frequency of Ki67^+^ of each of the Tr1 populations. (I) Representative flow cytometry of annexin V and 7AAD co-staining. Frequency of (J) 7AAD^-^AV^+^, (K) 7AAD^+^AV^-^, and (L) 7AAD^+^AV^+^ Tr1 cells. Each symbol is a different biological replicate, data shown as mean +/- SEM, (B-D) n=8, (F) n=8, and (J-L) n=9 biological replicates total from 2 independent experiments. Statistical analysis using one-way ANOVA with Bonferroni’s post-test *p<0.05, **p<0.01, ***p <0.001, ****p<0.0001.

P2RX7 is a surface purinergic receptor which senses extracellular ATP and can induce apoptosis (Di Virgilio et al., 2017). It has also been shown to regulate metabolic fitness in CD8^+^ T cells (Borges da Silva et al., 2018) but has not previously been implicated in Tr1 cell function. *P2rx7* was found to be most highly expressed by CD49b^+^ Tr1 populations (**Fig. S7**). In keeping with this, flow cytometric analysis revealed a significantly higher percentage of CD49b^+^ Tr1 cells were P2RX7^+^ compared to DP and LAG-3^+^ Tr1 cells (**Fig 7E, F**). Whether there were differences in fitness, proliferative capacity, and survival between the Tr1 cell populations was assessed by Ki67, Annexin V, and 7AAD staining. No differences were observed in cell cycling as reported by intranuclear Ki67 staining **(Fig 7G, H)**. However, when assessing fitness and death, DP and CD49b^+^ Tr1 cells contained a greater frequency of pre-apoptotic Annexin V^+^ cells compared to the other populations **(Fig 7I-L)**. In addition, the CD49b^+^ population contained the greatest proportion of 7AAD and Annexin V dual positive cells that had undergone apoptosis **(Fig 7I-L)**. These data are consistent with CD49b^+^ Tr1 cells undergoing a higher rate of apoptosis than the other Tr1 cell subsets and further suggests that CD49b^+^ Tr1 cells represent a more terminally differentiated Tr1 cell phenotype. Altogether, the transcriptional and functional data suggested a possible step-wise increase in T cell activation from DN to LAG-3^+^ to DP, with CD49b^+^ Tr1 cells representing an activated phenotype that had increased susceptibility to apoptosis.

### Plasticity of lung-derived Tr1 cells generated in response to IAV infection

To test whether an exclusively linear step-wise progression between the 4 Tr1 cell phenotypes observed in IAV infection exists, an adoptive transfer strategy was utilised. Each of the four Tr1 cell populations were FACS-sorted from lungs on day 7 post-infection (**Fig. S9**) and transferred into infection-matched WT hosts. Due to the technical challenges of working with small numbers of lung-derived Tr1 cells, despite pooling from multiple experiments, insufficient DN, DP, and CD49b^+^ Tr1 cells were recovered following transfer for robust analysis of their fate and plasticity. However, as the dominant population of Tr1 cells, the LAG-3^+^ Tr1 cells could be tracked reliably and were sufficient to test the hypothesis that Tr1 cells undergo a unidirectional linear progression through the observed phenotypes during acute IAV infection. On day 9 post-infection, tissues were collected for analysis of transferred cells **(Fig 8A-C)**. LAG-3^+^ Tr1 cells predominately maintained expression of IL-10 post-transfer and did not begin to express FOXP3 **(Fig 8D)**, indicating a degree of stability of the Tr1 cell phenotype, at least across this period of time. Regarding plasticity of LAG-3 and CD49b co-expression, LAG-3^+^ Tr1 cells either maintained their LAG-3^+^ phenotype or adopted either a DP or DN phenotype following adoptive transfer. Very few LAG-3^+^ Tr1 cells converted to a CD49b^+^ single-positive phenotype over the period of time analysed **(Fig 8D-F)**. Together, these experiments determined that LAG-3^+^ Tr1 cells exhibit stable IL-10 production but have the potential to rapidly adopt different expression profiles of LAG-3 and CD49b co-expression. Despite being limited to evaluation of transfer of the LAG-3^+^ Tr1 cells, these data establish that there is significant plasticity in Tr1 cell phenotypes during IAV infection and that there is not a uniform linear step-wise progression evident through Tr1 cell fates.

**Figure 8:**
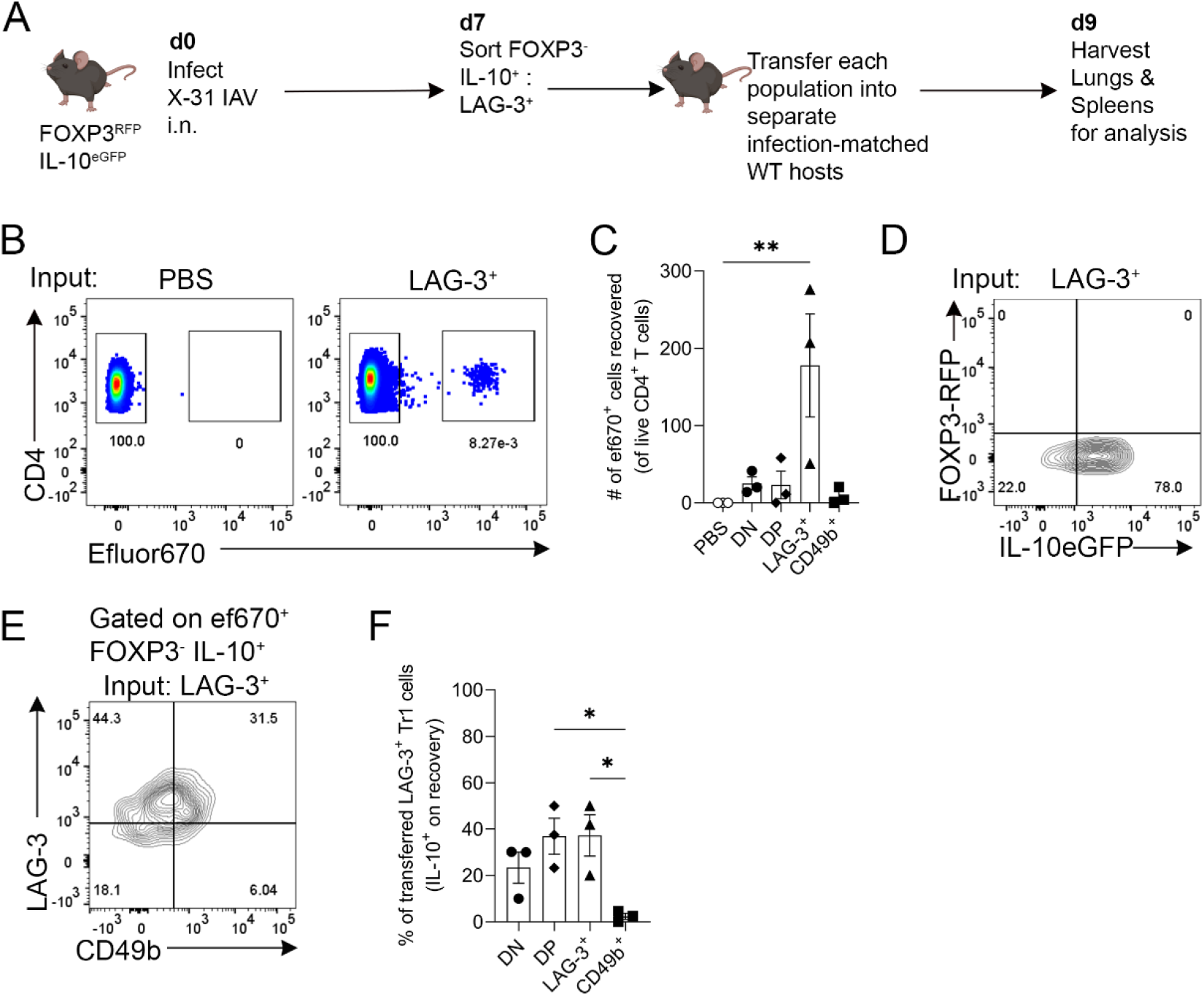
Plasticity of influenza-induced LAG-3^+^ Tr1 cells from the lung. (A) Experimental schematic. FOXP3^RFP^IL-10^GFP^ mice and age-matched C57BL/6 mice were infected with X-31 i.n. On day 7 post-infection lungs from 3-4 FOXP3^RFP^IL-10^GFP^ mice were pooled and Tr1 cells were FACS-sorted (Live, CD3^+^, CD4^+^, CD44^+^, FOXP3^-^, IL-10^+^) based on expression of LAG-3 and CD49b and transferred i.v. into separate infection-matched hosts. On day 9 post-infection the lungs and spleens were analysed for Tr1 cell presence and plasticity. (B) Concatenated flow cytometry of Tr1 cell detection (pooled from 3 independent experiments). (C) The number of ef670^+^ transferred cells recovered from lungs of recipients on day 9 post-infection. (D) Concatenated lungs and spleens from LAG-3^+^ recipient mice showing expression of FOXP3 and IL-10. (E) LAG-3 and CD49b expression by input LAG-3^+^ transferred cells that maintain IL-10^+^ (IL-10^+^). (F) The frequency of LAG-3 and CD49b expression by IL-10^+^ of transferred LAG-3^+^ Tr1 cells. Data are shown as mean +/- SEM, n=3 biological replicates from 3 independent experiments. Statistical analysis completed using one-way ANOVA with Bonferroni’s correction * p < 0.05, ** p < 0.01. (Figure made with Biorender.com)

## Discussion

Tr1 cells have been shown to play an important regulatory role by suppressing effector responses that limit chronic inflammation (Jankovic et al., 2007; Neumann et al., 2014; Roers et al., 2004). In this study we have shown for the first time that these cells also contribute to regulation of immune responses in the context of acute infection. These cells are the dominant T cell source of IL-10 in the lung of IAV-infected mice, greatly outnumbering IL-10^+^ Tregs, and in their absence there is delayed recovery from infection-induced weight loss, suggesting a role for Tr1 cells during the resolution of acute inflammation similar to that described in chronic inflammation (Gagliani et al., 2015). Intriguingly, we also reveal that Tr1 cells generated in response to IAV-infection have atypical surface phenotypes and exhibit heterogeneous expression of the Tr1 cell characteristic markers, LAG-3 and CD49b. Four distinct Tr1 cell populations which exhibited differences in their kinetics, phenotype, and function were observed. Using this model of Tr1 cell generation in acute infection we have revealed multiple novel insights into Tr1 cell biology, including new information on their development, heterogeneity, mechanisms of suppression, and phenotypic stability.

Some studies in humans and mice have indicated that co-expression of LAG-3 and CD49b in CD4^+^ T cells strongly enriches for Tr1 cells and these markers have been suggested to be used as reliable surrogates of a Tr1 cell phenotype (Brockmann et al., 2018; Huang et al., 2017; Gagliani et al., 2013). However, the general applicability of these markers has been questioned in other settings (Yao et al., 2015; Zhang et al., 2017), and has been shown to not be exclusively specific for Tr1 cells (Huang et al., 2018). The current study demonstrates that a LAG-3 CD49b co-expressing phenotype is not a general indicator of Tr1 cells. We clearly show that >75% of FOXP3^-^ IL-10^+^ T cells found in lungs of IAV-infected do not co-express LAG-3 and CD49b but meet all other defining criteria of Tr1 cells (i.e. are capable of suppressing bystander T cell proliferation and do not produce IL-4 or IL-17). The fact that DP Tr1 cells do not predominate in the acute resolving infection setting studied here is perhaps due to less potent T cell activation than occurs in other models previously studied (Brockmann et al., 2018; Gagliani et al., 2013; Huang et al., 2017). Our transcriptomic analysis indicated that DP Tr1 cells expressed more genes associated with T cell functionality and also had the highest level of TIM-3 expression, which may indicate that DP Tr1 cells represent a more tissue-infiltrated and functionally active stage of Tr1 cell biology than the other populations. Indeed, we found that a substantial fraction of LAG-3^+^ Tr1 cells converted to DP following transfer and also revealed that DP Tr1 cells were found deeper within the infected tissue (more parenchymal and localised in airways) than other subsets of Tr1 cells. Overall, the current study has established significant heterogeneity within the Tr1 cell compartment during acute infection. Although DP Tr1 cells were not the only population identified, they did appear to express the most genes associated with regulatory function, and hence may represent the most functional Tr1 cells.

One important feature of the different populations of Tr1 cells that emerged during acute IAV infection was differential dependence on IL-10 for suppression. Specifically, DP and CD49b^+^ cells utilised IL-10 but the other Tr1 cell phenotypes suppressed independently of IL-10. Moreover, neutralising IL-10 only partially inhibited suppression by DP and CD49b^+^ cells. Together, these observations indicate that other molecular mechanisms are used by those cells to exert suppression. While the mechanisms by which Tr1 cells exert suppression *in vivo* remain incompletely understood and vary from study-to-study, the most commonly reported mechanisms reported to be used by Tr1 cells to suppress effector T cell division are either IL-10 or via co-inhibitory receptors. For example, Tr1 cell-mediated suppression by Tr1 cells isolated from a model of colitis is dependent on IL-10 (Huber et al., 2011; Brockmann et al., 2017). However, post α-CD3 treatment, small intestinal Tr1 cells were enriched for expression of many co-inhibitory receptors and potently suppressed effector T cell proliferation *in vitro*, whereas splenic Tr1 cells from the same mice expressed fewer co-inhibitory receptors and were not as suppressive (Brockmann et al., 2018). In addition, Tr1 cells in models of antigen tolerization and allergy were shown to inhibit T cell proliferation via cell-contact mediated mechanisms dependent on PD-1 and CTLA-4 (Vieira et al., 2004; Akdis et al., 2004; Meiler et al., 2008). Studies of Tallo10 cells, a polarised human Tr1 cell clinical product, have revealed that Tr1 cells in this context suppress T cell proliferation via CTLA-4, PD-1, and IL-10 (Chen et al., 2021). Given the results from the current study, it will be of interest to determine any cell-contact dependent mechanisms of suppression mediated by Tr1 cell subpopulations isolated from IAV-infected lungs. Notably, DN Tr1 cells are suppressive independently of IL-10 and do not express LAG-3, which rules out a contribution of LAG-3 to their suppressive capacity. In addition, PD-1 and TIM-3 were highly expressed by all populations of Tr1 cells that developed in response to acute IAV infection, with particularly high expression of TIM-3 in the DP population, noting that in Tregs, TIM-3 can confer increased suppressive potential (Gautron et al., 2014). However, whether TIM-3 or other co-inhibitory receptors are actively involved in Tr1-mediated immune suppression in the context of acute IAV infection remains to be determined in future.

The circumstances under which Tr1 cells emerge, contribute to immune responses, and their subsequent fate remains somewhat enigmatic. Currently it is not definitively known whether these cells emerge *in vivo* directly from naïve precursors or adopt an IL-10-secreting phenotype after transitioning through another effector program, or can come from either pathway (Pot et al., 2009; Zhang et al., 2020; Gagliani et al., 2015; Huang et al., 2017). Analysis of the appearance of Tr1 cells in the present study indicate that, unlike Tregs, Tr1 cells do not substantially increase in frequency in draining lymph node or spleen following infection but accumulate abruptly in lungs on day 7. This indicates that it is likely that a substantial fraction of Tr1 cells derive from effector T cells that adopt IL-10 expression *in situ*. However, this will require future analysis of appropriate fate-reporter mice, such as has been shown for Th17 cells that transition to Tr1 in the context of autoimmunity (Gagliani et al., 2015). The kinetics of the appearance of Tr1 cells in the present study are quite consistent with previous studies in LCMV, *N*.*Brasiliensis* infection, and following lethal influenza infection (Huang et al., 2017; Gagliani et al., 2013), where these cells were also noted to appear between day 5 and day 7. However, the kinetics of the persistence of Tr1 cells in the present study substantially differs from that reported in chronic LCMV infection where Tr1 cells continued to accumulate up to day 20 (Parish et al., 2014), which is in contrast to the abrupt loss of Tr1 cells observed here by day 10 post-infection. This strict temporal control of Tr1 cells is suggestive of an important window for regulation of the anti-IAV immune response and raises important questions as to the fate and plasticity of these cells in this context.

While the data in this study demonstrate the generation and function of Tr1 cells in acute respiratory virus infection, the fate and persistence of Tr1 cells after response resolution were not fully resolved. While most Tr1 cells in the lung were lost between day 7 and day 10 post-IAV infection, it was notable that DN Tr1 cells persisted at day 10. On the surface, this initially suggested that DN Tr1 cells represented an endpoint of Tr1 cell fate. However, transcriptomic analysis instead suggested that DN Tr1 cells were the least differentiated Tr1 population (e.g. expressing higher levels of Tfh-like genes such as *Cxcr5* and *Id3*). Furthermore, the CD49b^+^ Tr1 cells were undergoing a higher rate of apoptosis compared to the other populations, suggesting that they represent an endpoint of Tr1 cell fate. An explanation for these findings is that the DN Tr1 cells observed in this study are a heterogeneous population of IL-10-secreting T cells that contains *bona fide* Tr1 cells that have not yet upregulated either CD49b or LAG-3 and also contains cells that have downregulated these markers as they transition to alternative T cell fates or undergo apoptosis. These questions could be addressed by developing genetic fate-reporting systems for Tr1 cells such as have been described for Th1, Th2, Treg and Th17 cells (Croxford et al., 2009; Lee et al., 2022; Rubtsov et al., 2010). In support of the concept that Tr1 cells exhibit plasticity *in vivo*, our data indicate that LAG-3^+^ Tr1 cells in acute IAV infection were capable of transitioning to non-Tr1 cells (loss of IL-10 expression) and also to DP and DN Tr1 cells. This suggests that there is substantial plasticity between the observed Tr1 cell phenotypes that emerge during acute IAV infection and that there is not a linear unidirectional transition between these phenotypes. Further investigation to untangle this complex web of cellular phenotypes using fate-mapping and single cell approaches is warranted to better understand these key regulatory cells in future.

## Conclusion

This study provides the first investigation of Tr1 cells in the context of an acute resolving model of infection, with a lack of Tr1 cells impeding resolution of IAV infection. Four distinct Tr1 cell phenotypes based on expression of LAG-3 and/or CD49b were observed in acute infection that exhibited different localisation, dependence on IL-10 to suppress T cell proliferation and had distinct transcriptional features. Our data support the concept of Tr1 cells contributing to resolution of acute respiratory infection. Thus, there is potential for Tr1 cells to be therapeutically exploited to regulate the adaptive immune response to limit tissue damage.

## Limitations of this study

The *Il27ra*^*-/-*^ mouse was used as a model of Tr1 deficiency. As this is a broad knockout mouse, other IL-27-dependent cell populations may also be affected. To further elucidate the extent of Tr1 contributions to regulation of immune responses other viruses and infection models should be investigated and these should also be examined in human patients during the course of acute respiratory infection.

## Acknowledgements

We would like to thank all staff at laboratory animal services at the University of Adelaide for assistance with animal husbandry, Dr. Randall Grose at South Australian Health and Medical Research Institute for assistance with cell sorting, Professor Stephen Turner (Monash University) for providing the X-31-OVA_323-339_ IAV, and Professor Christian Engwerda (QIMR) for providing IL-27ra^-/-^ mice.

## References

Agata, Y., A. Kawasaki, H. Nishimura, Y. Ishida, T. Tsubat, H. Yagita, and T. Honjo. 1996. Expression of the PD-1 antigen on the surface of stimulated mouse T and B lymphocytes. Int. Immunol. 8:765–772. doi:10.1093/intimm/8.5.765.

Akdis, M., J. Verhagen, A. Taylor, F. Karamloo, C. Karagiannidis, R. Crameri, S. Thunberg, G. Deniz, R. Valenta, H. Fiebig, C. Kegel, R. Disch, C.B. Schmidt-Weber, K. Blaser, and C.A. Akdis. 2004. Immune Responses in Healthy and Allergic Individuals Are Characterized by a Fine Balance between Allergen-specific T Regulatory 1 and T Helper 2 Cells. J. Exp. Med. 199:1567–1575. doi:10.1084/jem.20032058.

Anderson, K.G., K. Mayer-Barber, H. Sung, L. Beura, B.R. James, J.J. Taylor, L. Qunaj, T.S. Griffith, V. Vezys, D.L. Barber, and D. Masopust. 2014. Intravascular staining for discrimination of vascular and tissue leukocytes. Nat. Protoc. 9:209–222. doi:10.1038/nprot.2014.005.

Andrews, S. 2019. Babraham Bioinformatics - FastQC A Quality Control tool for High Throughput Sequence Data. Babraham Bioinforma.

Arpaia, N., J.A. Green, B. Moltedo, A. Arvey, S. Hemmers, S. Yuan, P.M. Treuting, and A.Y. Rudensky. 2015. A Distinct Function of Regulatory T Cells in Tissue Protection. Cell. 162:1078–1089. doi:10.1016/j.cell.2015.08.021.

Bacchetta, R., R. De, H. Yssel, and J. Abrams. 1990. Host-reactive CD4+ and CD8+ T cell clones isolated from a human chimera produce IL-5, IL-2, IFN-gamma and granulocyte/macrophage-colony-stimulating factor but not IL-4. J. Immunol. 144:902–908.

Batten, M., N.M. Kljavin, J. Li, M.J. Walter, F.J. de Sauvage, and N. Ghilardi. 2008. Cutting Edge: IL-27 Is a Potent Inducer of IL-10 but Not FoxP3 in Murine T Cells. J. Immunol. 180:2752–2756. doi:10.4049/jimmunol.180.5.2752.

Borges da Silva, H., L.K. Beura, H. Wang, E.A. Hanse, R. Gore, M.C. Scott, D.A. Walsh, K.E. Block, R. Fonseca, Y. Yan, K.L. Hippen, B.R. Blazar, D. Masopust, A. Kelekar, L. Vulchanova, K.A. Hogquist, and S.C. Jameson. 2018. The purinergic receptor P2RX7 directs metabolic fitness of long-lived memory CD8+ T cells. Nature. 559:264–268. doi:10.1038/s41586-018-0282-0.

Brockmann, L., N. Gagliani, B. Steglich, A.D. Giannou, J. Kempski, P. Pelczar, M. Geffken, B. Mfarrej, F. Huber, J. Herkel, Y.Y. Wan, E. Esplugues, M. Battaglia, C.F. Krebs, R.A. Flavell, and S. Huber. 2017. IL-10 Receptor Signaling Is Essential for T _R_1 Cell Function In Vivo. J. Immunol. 198:1130–1141. doi:10.4049/jimmunol.1601045.

Brockmann, L., S. Soukou, B. Steglich, P. Czarnewski, L. Zhao, S. Wende, T. Bedke, C. Ergen, C. Manthey, T. Agalioti, M. Geffken, O. Seiz, S.M. Parigi, C. Sorini, J. Geginat, K. Fujio, T. Jacobs, T. Roesch, J.R. Izbicki, A.W. Lohse, R.A. Flavell, C. Krebs, J.-A. Gustafsson, P. Antonson, M.G. Roncarolo, E.J. Villablanca, N. Gagliani, and S. Huber. 2018. Molecular and functional heterogeneity of IL-10-producing CD4+ T cells. Nat. Commun. 9:5457. doi:10.1038/s41467-018-07581-4.

Chen, P.P., A.-M. Cepika, R. Agarwal-Hashmi, G. Saini, M.J. Uyeda, D.M. Louis, B. Cieniewicz, M. Narula, L.C. Amaya Hernandez, N. Harre, L. Xu, B.C. Thomas, X. Ji, P. Shiraz, K.M. Tate, D. Margittai, N. Bhatia, E. Meyer, A. Bertaina, M.M. Davis, R. Bacchetta, and M.G. Roncarolo. 2021. Alloantigen-specific type 1 regulatory T cells suppress through CTLA-4 and PD-1 pathways and persist long-term in patients. Sci. Transl. Med. 13:eabf5264. doi:10.1126/scitranslmed.abf5264.

Collison, L.W., and D.A.A. Vignali. 2011. In Vitro Treg Suppression Assays. Methods Mol. Biol. Clifton NJ. 707:21–37. doi:10.1007/978-1-61737-979-6_2.

Croxford, A.L., F.C. Kurschus, and A. Waisman. 2009. Cutting Edge: An IL-17F-CreEYFP Reporter Mouse Allows Fate Mapping of Th17 Cells. J. Immunol. 182:1237–1241. doi:10.4049/jimmunol.182.3.1237.

Di Virgilio, F., D. Dal Ben, A.C. Sarti, A.L. Giuliani, and S. Falzoni. 2017. The P2X7 Receptor in Infection and Inflammation. Immunity. 47:15–31. doi:10.1016/j.immuni.2017.06.020.

Dobin, A., C.A. Davis, F. Schlesinger, J. Drenkow, C. Zaleski, S. Jha, P. Batut, M. Chaisson, and T.R. Gingeras. 2013. STAR: ultrafast universal RNA-seq aligner. Bioinformatics. 29:15–21. doi:10.1093/bioinformatics/bts635.

Fulton, R.B., D.K. Meyerholz, and S.M. Varga. 2010. Foxp3+ CD4 Regulatory T Cells Limit Pulmonary Immunopathology by Modulating the CD8 T Cell Response during Respiratory Syncytial Virus Infection. J. Immunol. 185:2382–2392. doi:10.4049/jimmunol.1000423.

Gagliani, N., C.F. Magnani, S. Huber, M.E. Gianolini, M. Pala, P. Licona-Limon, B. Guo, D.R. Herbert, A. Bulfone, F. Trentini, C. Di Serio, R. Bacchetta, M. Andreani, L. Brockmann, S. Gregori, R.A. Flavell, and M.-G. Roncarolo. 2013. Coexpression of CD49b and LAG-3 identifies human and mouse T regulatory type 1 cells. Nat. Med. 19:739–746. doi:10.1038/nm.3179.

Gagliani, N., M.C.A. Vesely, A. Iseppon, L. Brockmann, H. Xu, N.W. Palm, M.R. de Zoete, P. Licona-Limón, R.S. Paiva, T. Ching, C. Weaver, X. Zi, X. Pan, R. Fan, L.X. Garmire, M.J. Cotton, Y. Drier, B. Bernstein, J. Geginat, B. Stockinger, E. Esplugues, S. Huber, and R.A. Flavell. 2015. TH17 cells transdifferentiate into regulatory T cells during resolution of inflammation. Nature. 523:221–225. doi:10.1038/nature14452.

Gautron, A.-S., M. Dominguez-Villar, M. de Marcken, and D.A. Hafler. 2014. Enhanced suppressor function of TIM-3+FoxP3+ regulatory T cells. Eur. J. Immunol. 44:2703–2711. doi:10.1002/eji.201344392.

Ghalib, H.W., M.R. Piuvezam, Y.A. Skeiky, M. Siddig, F.A. Hashim, A.M. el-Hassan, D.M. Russo, and S.G. Reed. 1993. Interleukin 10 production correlates with pathology in human Leishmania donovani infections. J. Clin. Invest. 92:324–329. doi:10.1172/JCI116570.

Gregori, S., D. Tomasoni, V. Pacciani, M. Scirpoli, M. Battaglia, C.F. Magnani, E. Hauben, and M.-G. Roncarolo. 2010. Differentiation of type 1 T regulatory cells (Tr1) by tolerogenic DC-10 requires the IL-10–dependent ILT4/HLA-G pathway. Blood. 116:935–944. doi:10.1182/blood-2009-07-234872.

Groux, H., A. O’Garra, M. Bigler, M. Rouleau, S. Antonenko, J.E. de Vries, and M.G. Roncarolo. 1997. A CD4+T-cell subset inhibits antigen-specific T-cell responses and prevents colitis. Nature. 389:737–742. doi:10.1038/39614.

Hagenbaugh, A., S. Sharma, S.M. Dubinett, S.H.-Y. Wei, R. Aranda, H. Cheroutre, D.J. Fowell, S. Binder, B. Tsao, R.M. Locksley, K.W. Moore, and M. Kronenberg. 1997. Altered Immune Responses in Interleukin 10 Transgenic Mice. J. Exp. Med. 185:2101–2110.

Hastings, W.D., D.E. Anderson, N. Kassam, K. Koguchi, E.A. Greenfield, S.C. Kent, X.X. Zheng, T.B. Strom, D.A. Hafler, and V.K. Kuchroo. 2009. TIM-3 is expressed on activated human CD4+ T cells and regulates Th1 and Th17 cytokines. Eur. J. Immunol. 39:2492–2501. doi:10.1002/eji.200939274.

Heinzel, F.P., M.D. Sadick, S.S. Mutha, and R.M. Locksley. 1991. Production of interferon gamma, interleukin 2, interleukin 4, and interleukin 10 by CD4+ lymphocytes in vivo during healing and progressive murine leishmaniasis. Proc. Natl. Acad. Sci. 88:7011–7015. doi:10.1073/pnas.88.16.7011.

Hoffman, G.E., and E.E. Schadt. 2016. variancePartition: interpreting drivers of variation in complex gene expression studies. BMC Bioinformatics. 17:483. doi:10.1186/s12859-016-1323-z.

Holaday, B.J., M.M. Pompeu, S. Jeronimo, M.J. Texeira, A. de A. Sousa, A.W. Vasconcelos, R.D. Pearson, J.S. Abrams, and R.M. Locksley. 1993. Potential role for interleukin-10 in the immunosuppression associated with kala azar. J. Clin. Invest. 92:2626–2632.

Huang, W., S. Solouki, C. Carter, S.-G. Zheng, and A. August. 2018. Beyond Type 1 Regulatory T Cells: Co-expression of LAG3 and CD49b in IL-10-Producing T Cell Lineages. Front. Immunol. 9:2625. doi:10.3389/fimmu.2018.02625.

Huang, W., S. Solouki, N. Koylass, S.-G. Zheng, and A. August. 2017. ITK signalling via the Ras/IRF4 pathway regulates the development and function of Tr1 cells. Nat. Commun. 8:15871. doi:10.1038/ncomms15871.

Huber, S., N. Gagliani, E. Esplugues, W. O’Connor, F.J. Huber, A. Chaudhry, M. Kamanaka, Y. Kobayashi, C.J. Booth, A.Y. Rudensky, M.G. Roncarolo, M. Battaglia, and R.A. Flavell. 2011. Th17 Cells Express Interleukin-10 Receptor and Are Controlled by Foxp3− and Foxp3+ Regulatory CD4+ T Cells in an Interleukin-10-Dependent Manner. Immunity. 34:554–565. doi:10.1016/j.immuni.2011.01.020.

Jankovic, D., M.C. Kullberg, C.G. Feng, R.S. Goldszmid, C.M. Collazo, M. Wilson, T.A. Wynn, M. Kamanaka, R.A. Flavell, and A. Sher. 2007. Conventional T-bet+Foxp3− Th1 cells are the major source of host-protective regulatory IL-10 during intracellular protozoan infection. J. Exp. Med. 204:273–283. doi:10.1084/jem.20062175.

Kamanaka, M., S.T. Kim, Y.Y. Wan, F.S. Sutterwala, M. Lara-Tejero, J.E. Galán, E. Harhaj, and R.A. Flavell. 2006. Expression of Interleukin-10 in Intestinal Lymphocytes Detected by an Interleukin-10 Reporter Knockin tiger Mouse. Immunity. 25:941–952. doi:10.1016/j.immuni.2006.09.013.

Karwacz, K., E.R. Miraldi, M. Pokrovskii, A. Madi, N. Yosef, I. Wortman, X. Chen, A. Watters, N. Carriero, A. Awasthi, A. Regev, R. Bonneau, D. Littman, and V.K. Kuchroo. 2017. Critical role of IRF1 and BATF in forming chromatin landscape during type 1 regulatory cell differentiation. Nat. Immunol. 18:412–421. doi:10.1038/ni.3683.

Kolde, R. 2012. Pheatmap: pretty heatmaps. R Package Version. 1:726.

Law, C.W., Y. Chen, W. Shi, and G.K. Smyth. 2014. voom: precision weights unlock linear model analysis tools for RNA-seq read counts. Genome Biol. 15:R29. doi:10.1186/gb-2014-15-2-r29.

Lee, S.E., B.D. Rudd, and N.L. Smith. 2022. Fate-mapping mice: new tools and technology for immune discovery. Trends Immunol. 43:195–209. doi:10.1016/j.it.2022.01.004.

Levings, M.K., R. Sangregorio, and M.-G. Roncarolo. 2001. Human Cd25+Cd4+ T Regulatory Cells Suppress Naive and Memory T Cell Proliferation and Can Be Expanded in Vitro without Loss of Function. J. Exp. Med. 193:1295–1302. doi:10.1084/jem.193.11.1295.

Liu, F.D.M., E.E. Kenngott, M.F. Schröter, A. Kühl, S. Jennrich, R. Watzlawick, U. Hoffmann, T. Wolff, S. Norley, A. Scheffold, J.S. Stumhofer, C.J.M. Saris, J.M. Schwab, C.A. Hunter, G.F. Debes, and A. Hamann. 2014. Timed Action of IL-27 Protects from Immunopathology while Preserving Defense in Influenza. PLOS Pathog. 10:e1004110. doi:10.1371/journal.ppat.1004110.

Lu, C., D. Zanker, P. Lock, X. Jiang, J. Deng, M. Duan, C. Liu, P. Faou, M.J. Hickey, and W. Chen. 2019. Memory regulatory T cells home to the lung and control influenza A virus infection. Immunol. Cell Biol. 97:774–786. doi:10.1111/imcb.12271.

Lund, J.M., L. Hsing, T.T. Pham, and A.Y. Rudensky. 2008. Coordination of Early Protective Immunity to Viral Infection by Regulatory T Cells. Science. doi:10.1126/science.1155209.

Maintainer, B.P., M. Morgan, M. Carlson, D. Tenenbaum, S. Arora, V. Oberchain, K. Morrell, and L. Shepherd. 2022. AnnotationHub: Client to access AnnotationHub resources. doi:10.18129/B9.bioc.AnnotationHub.

Meiler, F., J. Zumkehr, S. Klunker, B. Rückert, C.A. Akdis, and M. Akdis. 2008. In vivo switch to IL-10– secreting T regulatory cells in high dose allergen exposure. J. Exp. Med. 205:2887–2898. doi:10.1084/jem.20080193.

Neumann, C., F. Heinrich, K. Neumann, V. Junghans, M.-F. Mashreghi, J. Ahlers, M. Janke, C. Rudolph, N. Mockel-Tenbrinck, A.A. Kühl, M.M. Heimesaat, C. Esser, S.-H. Im, A. Radbruch, S. Rutz, and A. Scheffold. 2014. Role of Blimp-1 in programing Th effector cells into IL-10 producers. J. Exp. Med. 211:1807–1819. doi:10.1084/jem.20131548.

Oca, M.M. de, R. Kumar, F. de L. Rivera, F.H. Amante, M. Sheel, R.J. Faleiro, P.T. Bunn, S.E. Best, L. Beattie, S.S. Ng, C.L. Edwards, W. Muller, E. Cretney, S.L. Nutt, M.J. Smyth, A. Haque, G.R. Hill, S. Sundar, A. Kallies, and C.R. Engwerda. 2016. Blimp-1-Dependent IL-10 Production by Tr1 Cells Regulates TNF-Mediated Tissue Pathology. PLOS Pathog. 12:e1005398. doi:10.1371/journal.ppat.1005398.

Parish, I.A., H.D. Marshall, M.M. Staron, P.A. Lang, A. Brüstle, J.H. Chen, W. Cui, Y.-C. Tsui, C. Perry, B.J. Laidlaw, P.S. Ohashi, C.T. Weaver, and S.M. Kaech. 2014. Chronic viral infection promotes sustained Th1-derived immunoregulatory IL-10 via BLIMP-1. J. Clin. Invest. 124:3455–3468. doi:10.1172/JCI66108.

Pedersen, H.W., Danielle Navarro, and Thomas Lin. ggplot2 Elegant Graphics for Data Analysis. 3rd ed. Springer.

Pot, C., H. Jin, A. Awasthi, S.M. Liu, C.-Y. Lai, R. Madan, A.H. Sharpe, C.L. Karp, S.-C. Miaw, I.-C. Ho, and V.K. Kuchroo. 2009. Cutting Edge: IL-27 Induces the Transcription Factor c-Maf, Cytokine IL-21, and the Costimulatory Receptor ICOS that Coordinately Act Together to Promote Differentiation of IL-10-Producing Tr1 Cells. J. Immunol. 183:797–801. doi:10.4049/jimmunol.0901233.

Reed, S.G., C.E. Brownell, D.M. Russo, J.S. Silva, K.H. Grabstein, and P.J. Morrissey. 1994. IL-10 mediates susceptibility to Trypanosoma cruzi infection. J. Immunol. 153:3135–3140.

Reiner, S.L., S. Zheng, Z.E. Wang, L. Stowring, and R.M. Locksley. 1994. Leishmania promastigotes evade interleukin 12 (IL-12) induction by macrophages and stimulate a broad range of cytokines from CD4+ T cells during initiation of infection. J. Exp. Med. 179:447–456. doi:10.1084/jem.179.2.447.

Ritchie, M.E., B. Phipson, D. Wu, Y. Hu, C.W. Law, W. Shi, and G.K. Smyth. 2015. limma powers differential expression analyses for RNA-sequencing and microarray studies. Nucleic Acids Res. 43:e47. doi:10.1093/nar/gkv007.

Roers, A., L. Siewe, E. Strittmatter, M. Deckert, D. Schlüter, W. Stenzel, A.D. Gruber, T. Krieg, K. Rajewsky, and W. Müller. 2004. T Cell–specific Inactivation of the Interleukin 10 Gene in Mice Results in Enhanced T Cell Responses but Normal Innate Responses to Lipopolysaccharide or Skin Irritation. J. Exp. Med. 200:1289–1297. doi:10.1084/jem.20041789.

Rogers, M.C., K.D. Lamens, N. Shafagati, M. Johnson, T.D. Oury, S. Joyce, and J.V. Williams. 2018. CD4 ^+^ Regulatory T Cells Exert Differential Functions during Early and Late Stages of the Immune Response to Respiratory Viruses. J. Immunol. 201:1253–1266. doi:10.4049/jimmunol.1800096.

Roncarolo, M.G., S. Gregori, R. Bacchetta, M. Battaglia, and N. Gagliani. 2018. The Biology of T Regulatory Type 1 Cells and Their Therapeutic Application in Immune-Mediated Diseases. Immunity. 49:1004–1019. doi:10.1016/j.immuni.2018.12.001.

Roncarolo, M.G., H. Yssel, J.L. Touraine, H. Betuel, J.E. De Vries, and H. Spits. 1988. Autoreactive T cell clones specific for class I and class II HLA antigens isolated from a human chimera. J. Exp. Med. 167:1523–1534. doi:10.1084/jem.167.5.1523.

Rubtsov, Y.P., R.E. Niec, S. Josefowicz, L. Li, J. Darce, D. Mathis, C. Benoist, and A.Y. Rudensky. 2010. Stability of the Regulatory T Cell Lineage in Vivo. Science. 329:1667–1671. doi:10.1126/science.1191996.

Ruckwardt, T.J., K.L. Bonaparte, M.C. Nason, and B.S. Graham. 2009. Regulatory T Cells Promote Early Influx of CD8+ T Cells in the Lungs of Respiratory Syncytial Virus-Infected Mice and Diminish Immunodominance Disparities. J. Virol. doi:10.1128/JVI.00036-09.

Schmittgen, T.D., and K.J. Livak. 2008. Analyzing real-time PCR data by the comparative CT method. Nat. Protoc. 3:1101–1108. doi:10.1038/nprot.2008.73.

Schubert, M., S. Lindgreen, and L. Orlando. 2016. AdapterRemoval v2: rapid adapter trimming, identification, and read merging. BMC Res. Notes. 9:88. doi:10.1186/s13104-016-1900-2.

Sehrawat, S., S. Suvas, P.P. Sarangi, A. Suryawanshi, and B.T. Rouse. 2008. In Vitro-Generated Antigen-Specific CD4+ CD25+ Foxp3+ Regulatory T Cells Control the Severity of Herpes Simplex Virus-Induced Ocular Immunoinflammatory Lesions. J. Virol. doi:10.1128/JVI.00697-08.

Vieira, P.L., J.R. Christensen, S. Minaee, E.J. O’Neill, F.J. Barrat, A. Boonstra, T. Barthlott, B. Stockinger, D.C. Wraith, and A. O’Garra. 2004. IL-10-Secreting Regulatory T Cells Do Not Express Foxp3 but Have Comparable Regulatory Function to Naturally Occurring CD4+CD25+ Regulatory T Cells. J. Immunol. 172:5986–5993. doi:10.4049/jimmunol.172.10.5986.

Wan, Y.Y., and R.A. Flavell. 2005. Identifying Foxp3-expressing suppressor T cells with a bicistronic reporter. Proc. Natl. Acad. Sci. U. S. A. 102:5126–5131. doi:10.1073/pnas.0501701102.

Ward, C.M., T.-H. To, and S.M. Pederson. 2020. ngsReports: a Bioconductor package for managing FastQC reports and other NGS related log files. Bioinformatics. 36:2587–2588. doi:10.1093/bioinformatics/btz937.

Ward, S.T., K.-K. Li, and S.M. Curbishley. 2014. A method for conducting suppression assays using small numbers of tissue-isolated regulatory T cells. MethodsX. 1:168–174. doi:10.1016/j.mex.2014.08.012.

Yao, Y., J. Vent-Schmidt, M.D. McGeough, M. Wong, H.M. Hoffman, T.S. Steiner, and M.K. Levings. 2015. Tr1 Cells, but Not Foxp3+ Regulatory T Cells, Suppress NLRP3 Inflammasome Activation via an IL-10–Dependent Mechanism. J. Immunol. 195:488–497. doi:10.4049/jimmunol.1403225.

Yu, G., L.-G. Wang, Y. Han, and Q.-Y. He. 2012. clusterProfiler: an R Package for Comparing Biological Themes Among Gene Clusters. OMICS J. Integr. Biol. 16:284–287. doi:10.1089/omi.2011.0118.

Yu, H., N. Gagliani, H. Ishigame, S. Huber, S. Zhu, E. Esplugues, K.C. Herold, L. Wen, and R.A. Flavell. 2017. Intestinal type 1 regulatory T cells migrate to periphery to suppress diabetogenic T cells and prevent diabetes development. Proc. Natl. Acad. Sci. 114:10443–10448. doi:10.1073/pnas.1705599114.

Zhang, H., A. Madi, N. Yosef, N. Chihara, A. Awasthi, C. Pot, C. Lambden, A. Srivastava, P.R. Burkett, J. Nyman, E. Christian, Y. Etminan, A. Lee, H. Stroh, J. Xia, K. Karwacz, P.I. Thakore, N. Acharya, A. Schnell, C. Wang, L. Apetoh, O. Rozenblatt-Rosen, A.C. Anderson, A. Regev, and V.K. Kuchroo. 2020. An IL-27-Driven Transcriptional Network Identifies Regulators of IL-10 Expression across T Helper Cell Subsets. Cell Rep. 33:108433. doi:10.1016/j.celrep.2020.108433.

Zhang, P., J.S. Lee, K.H. Gartlan, I.S. Schuster, I. Comerford, A. Varelias, M.A. Ullah, S. Vuckovic, M. Koyama, R.D. Kuns, K.R. Locke, K.J. Beckett, S.D. Olver, L.D. Samson, M. Montes de Oca, F. de Labastida Rivera, A.D. Clouston, G.T. Belz, B.R. Blazar, K.P. MacDonald, S.R. McColl, R. Thomas, C.R. Engwerda, M.A. Degli-Esposti, A. Kallies, S.-K. Tey, and G.R. Hill. 2017. Eomesodermin promotes the development of type 1 regulatory T (TR1) cells. Sci. Immunol. 2:eaah7152. doi:10.1126/sciimmunol.aah7152.

Zou, Q., B. Wu, J. Xue, X. Fan, C. Feng, S. Geng, M. Wang, and B. Wang. 2014. CD8+ Treg cells suppress CD8+ T cell-responses by IL-10-dependent mechanism during H5N1 influenza virus infection. Eur. J. Immunol. 44:103–114. doi:10.1002/eji.201343583.

