## Supplementary figures for "Novel populations of Tr1 cells contribute to the resolution of acute influenza A virus infection"

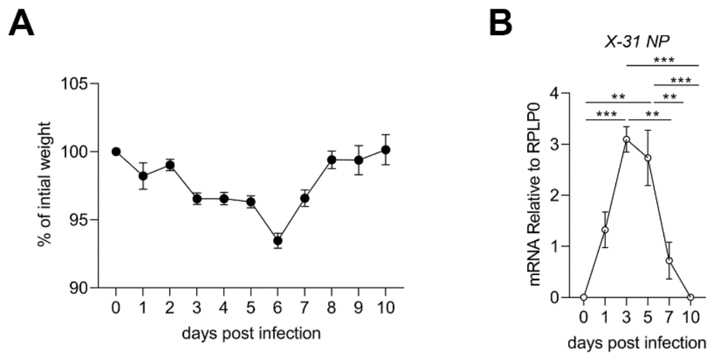


**Figure S1: Acute IAV infection model.**

(A) Average percentage of initial weight following i.n X-31 IAV infection of B6 mice. (B) Lungs were harvested on day 0 (naive) or days 1, 3, 5, 7 and 10 post-infection, RNA extracted, and viral load was assessed by RT-qPCR for X-31 nucleoprotein (NP) relative to housekeeping gene *(Rplp0)*. The symbols represent the mean +/- SEM, n=3-5 biological replicates total per time point (A), and n=50 biological replicates (B). Statistical analysis using one-way ANOVA with Bonferroni's post-test *p<0.05, **p<0.01, ***p <0.001, ****p<0.0001.

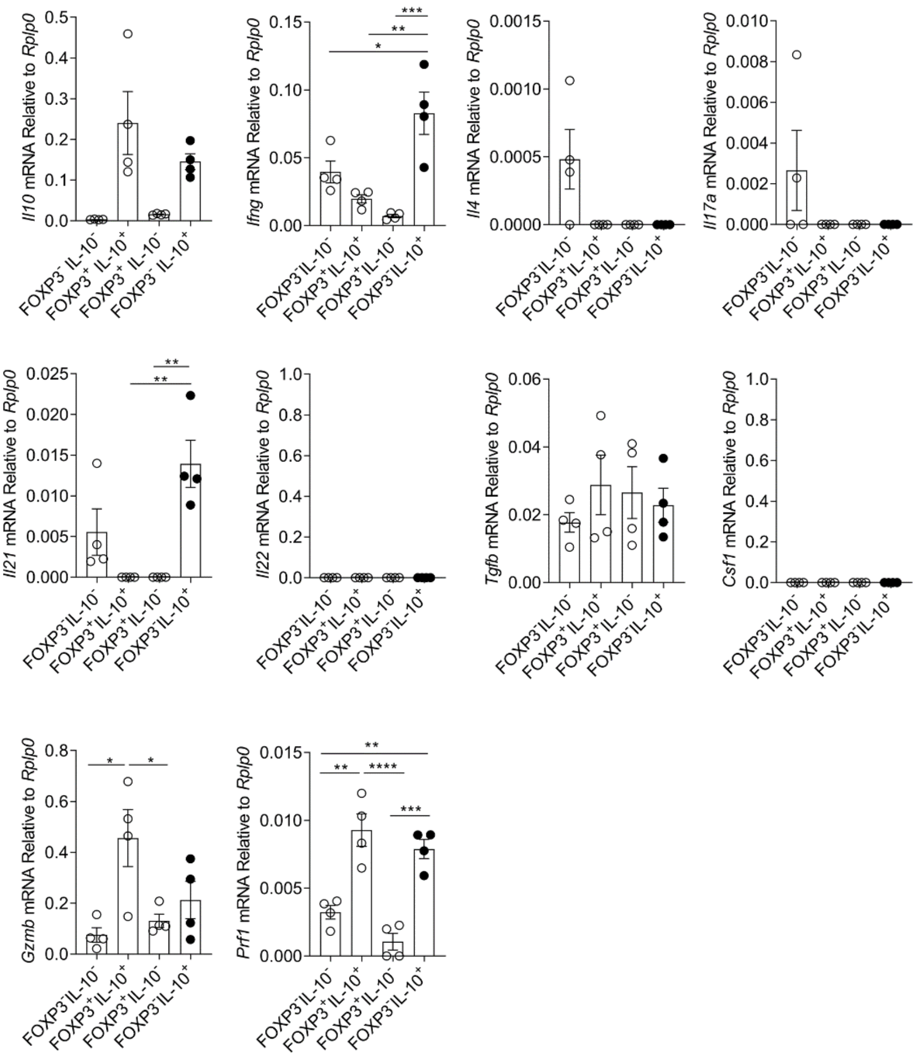


**Figure S2: Cytokines expressed by Tr1 cells.**

On day 7 post-infection CD4$^+$ T cell populations were sorted from lungs on day 7 post-infection based on FOXP3 and IL-10 expression. Cytokine expression by each of the populations was assessed by RT-qPCR relative to housekeeping gene *(Rplp0*). The symbols represent individual biological replicates +/- SEM, n=4 biological replicates total from 2 independent experiments. Statistical analysis using one-way ANOVA with Bonferroni's post-test *p<0.05, **p<0.01, ***p <0.001, ****p<0.0001.

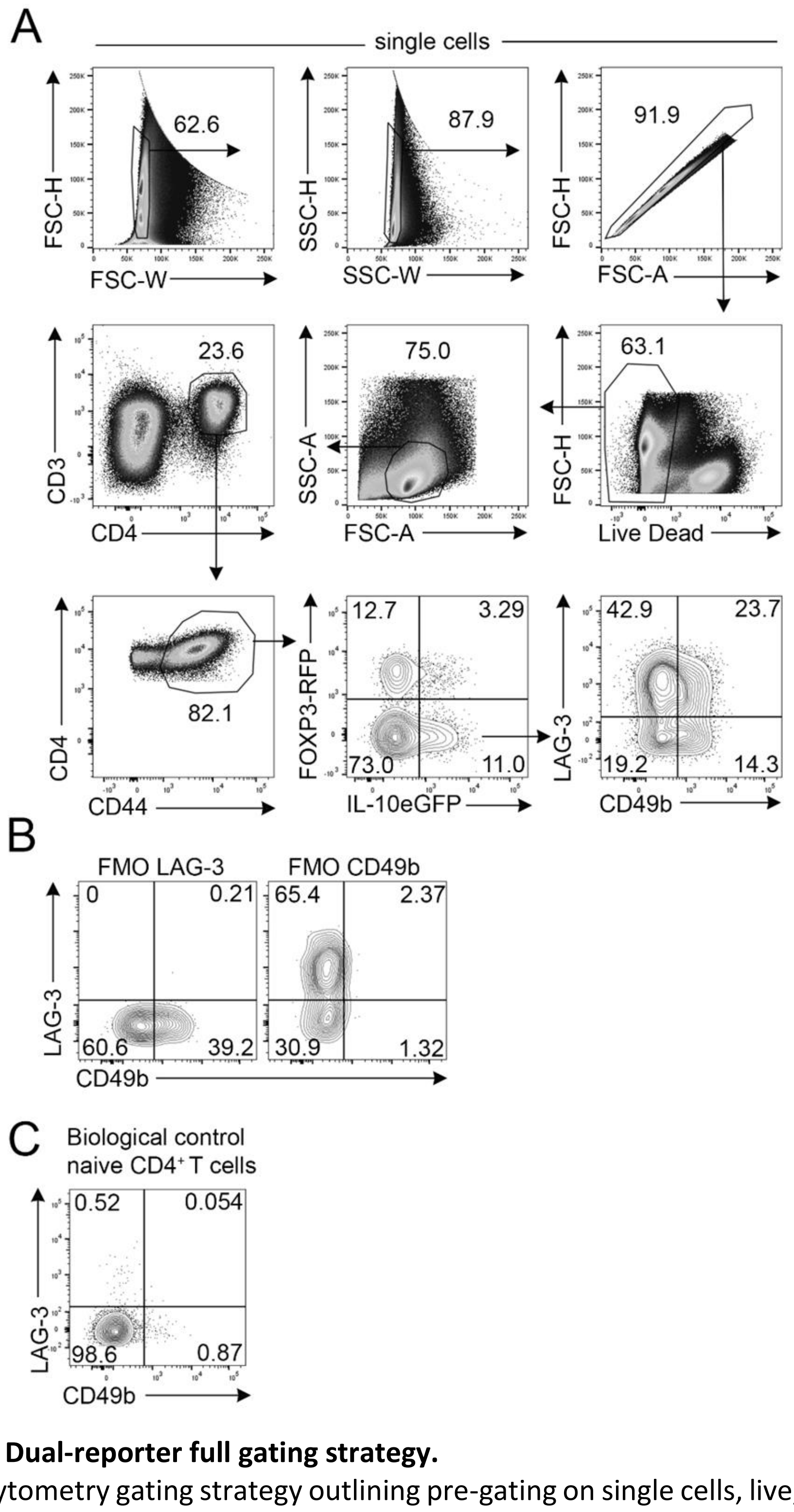


**Figure S3: Dual-reporter full gating strategy.**

(A) Flow cytometry gating strategy outlining pre-gating on single cells, live, lymphocytes, $CD3^+$ $CD4^+$ T cells prior to activated ($CD44^+$), $FOXP3^+$ $IL\text{-}10^+$ Tr1 cells. LAG-3 and CD49b populations are defined with a quadrant gate. (B) Fluorescence minus one (FMO) controls for LAG-3 and CD49b are used in combination with (C) Biological negative control for LAG-3 and CD49b expression (naive $CD4^+$ T cells).

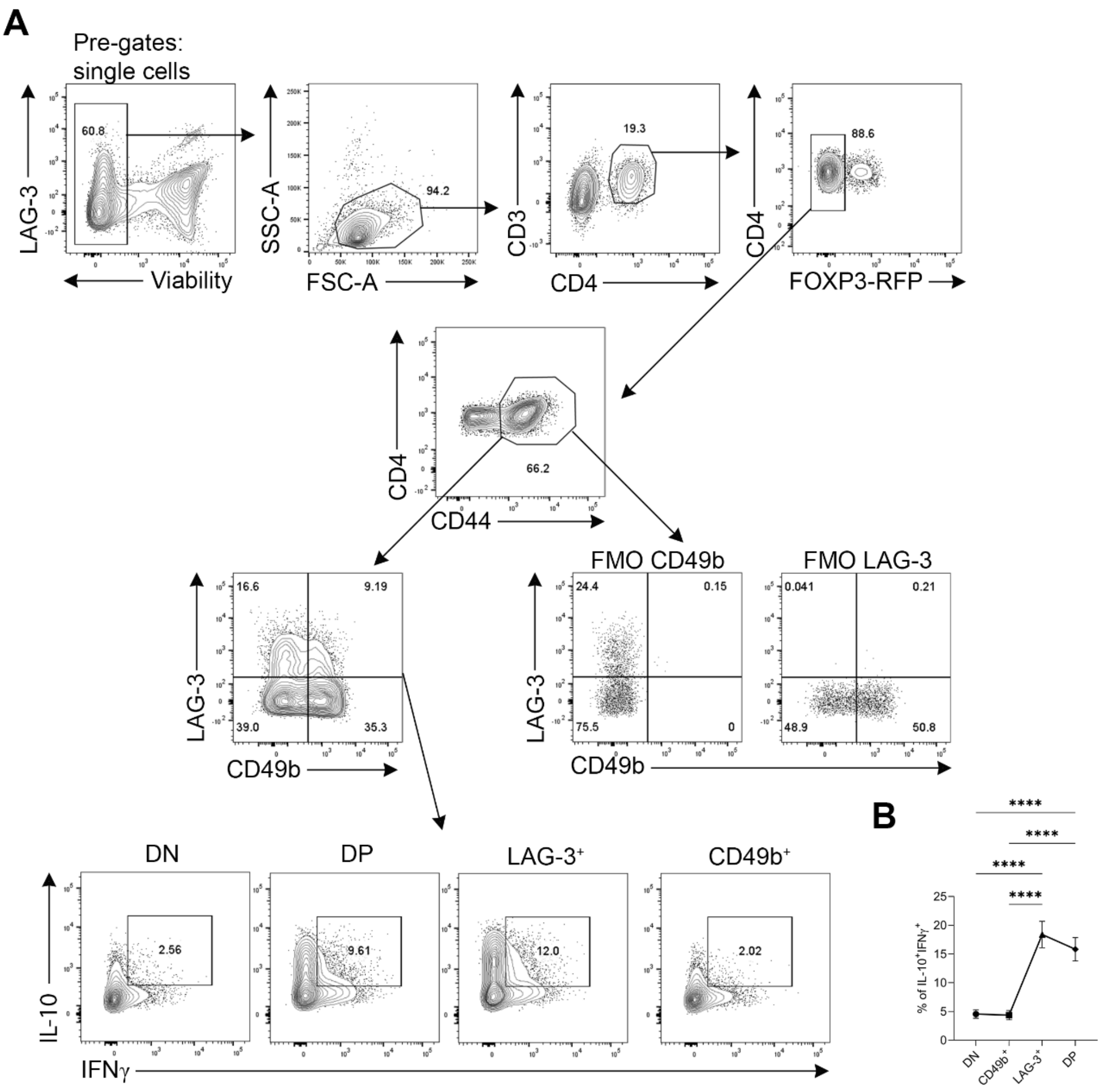


**Figure S4: Alternate dual-reporter full gating strategy identifying Tr1 cells based on IL-10 and IFNγ.**

(A) Alternate flow cytometry gating strategy outlining pre-gating on single cells, live, lymphocytes, $CD3^+$ $CD4^+$ T cells prior to $FOXP3^-$, activated ($CD44^+$), quadrant gates were then applied for LAG-3 and CD49b (FMO controls shown to the right). IL-10 and IFNγ co-expression was gated to compare relative enrichment within the four LAG-3 and CD49b populations. (B) Frequency of IL-10 and IFNγ co-expression by the four LAG-3 and CD49b populations. Mean is shown +/- SEM, n=12 biological replicates from 2 independent experiments. Statistical analysis using one-way ANOVA with Bonferroni's post-test *p<0.05, **p<0.01, ***p <0.001, ****p<0.0001.

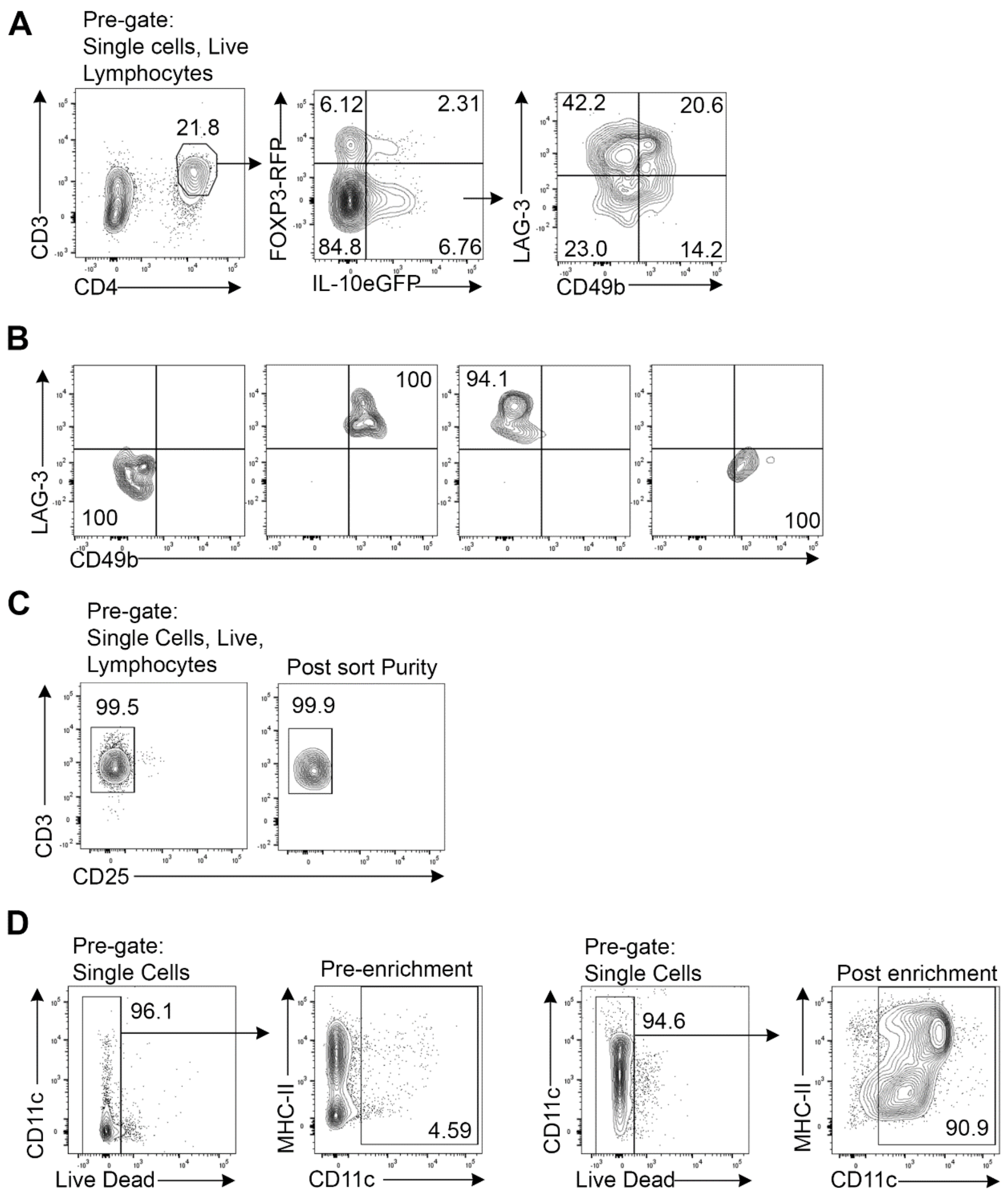


**Figure S5: Cell sorting and enrichment for suppression assays.**

Dual-reporter mice were infected with X-31 IAV. (A) Representative flow cytometry gating strategy for FACS-sorting FOXP3$^-$ IL-10$^+$ CD4$^+$ DN, DP, LAG-3$^+$, and CD49b$^+$ Tr1 cell populations for suppression assays. (B) Representative flow cytometry showing purity of the populations post sorting. (C) Representative pre- and post-sort purity of CD3$^+$CD25$^-$ effector T cells from SLOs post pan-CD3$^+$ T cell negative selection kit (StemCell). (D) Representative flow cytometry showing enrichment of CD11c$^+$ cells post CD11c positive selection kit (StemCell).

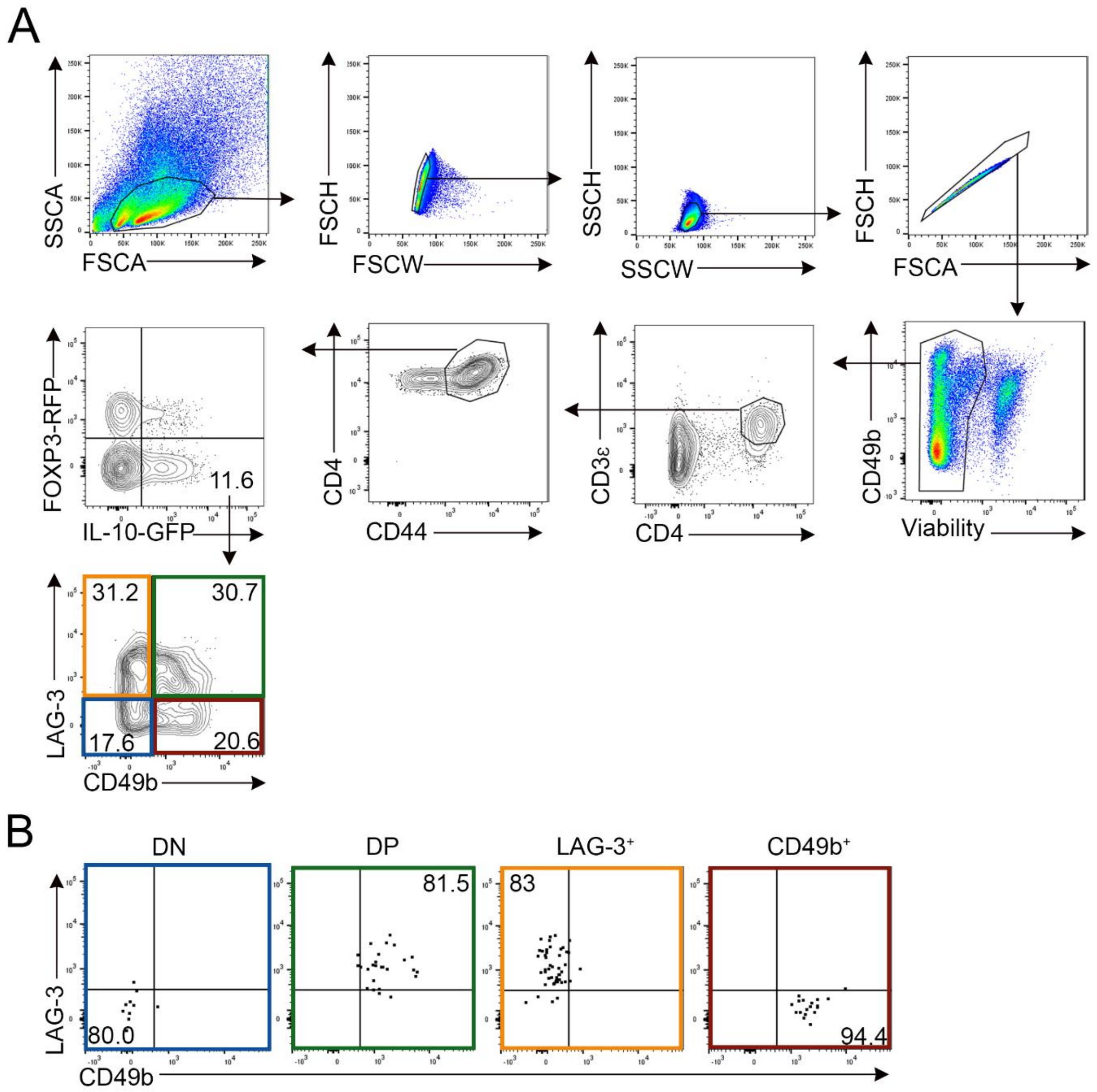


**Figure S6: Tr1 cell sort for sequencing.**

Dual-reporter mice were infected with X-31 IAV. (A) Representative flow cytometry gating strategy for FACS-sorting FOXP3$^-$ IL-10$^+$ CD4$^+$ DN, DP, LAG-3$^+$, and CD49b$^+$ Tr1 cell populations for RNA-sequencing. (B) Representative plots showing purity of the populations post sorting. Initially, the four Tr1 cell populations were sorted from n=6 biological replicates, in 2 independent experiments, after RNA preparation and QC analysis was conducted two biological replicates were excluded due to lower RNA quality.

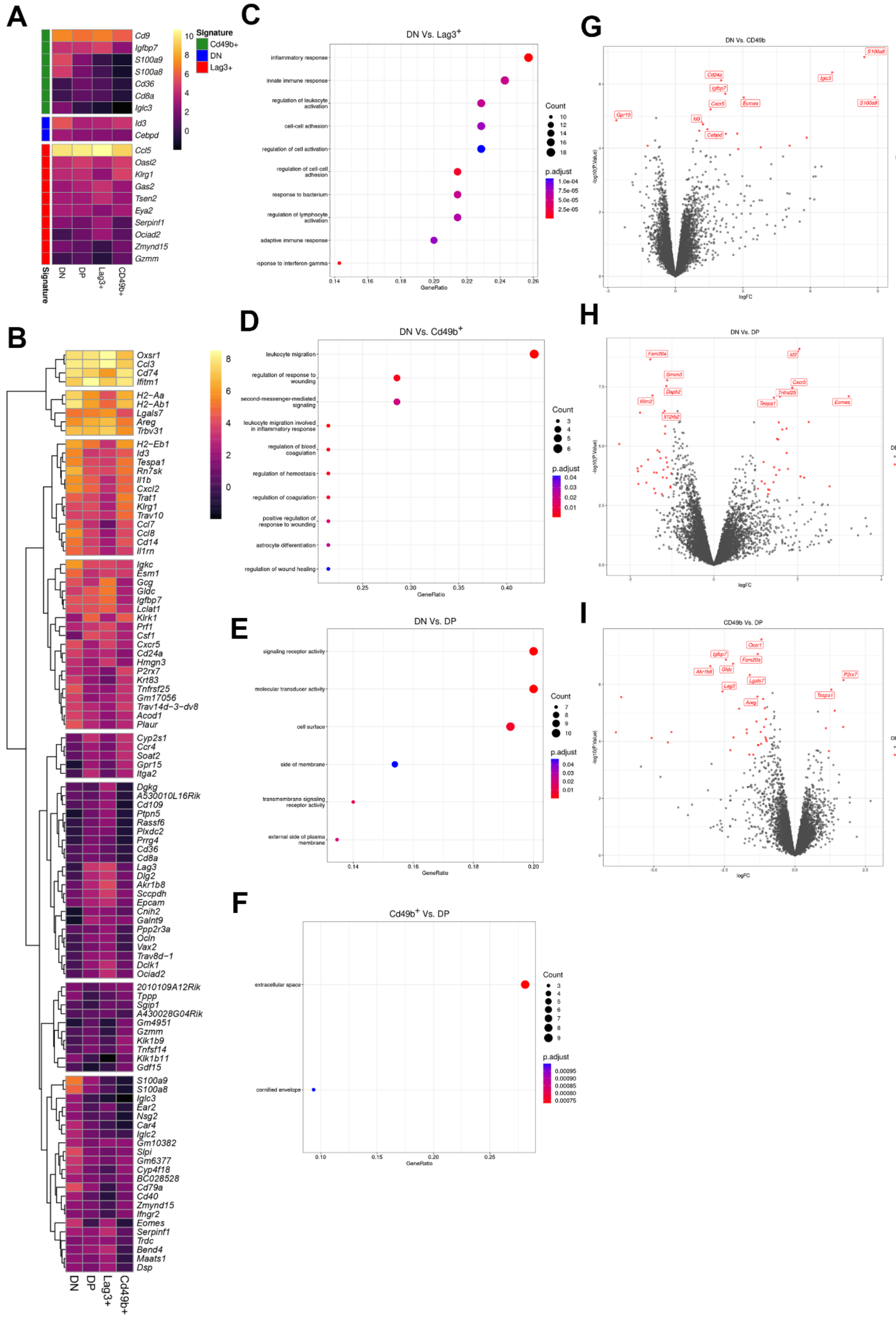


**Figure S7: Tr1 population comparisons.**

(A) Signature genes heatmap as defined by UpSet analysis, expressed as average logCPM. No significant unique signature was defined for the DP population. (B) Heatmap of the top 100 most DE genes ranked by LFC, with the scale indicating average logCPM within each cell type.

All genes were considered as DE in at least one comparison using an FDR-adjusted p-value < 0.05 and estimated logFC beyond the range ±1. Pathway analysis was conducted using clusterprofileR in R studio and dot plots are shown for the comparisons between (C) DN vs LAG-3$^{+}$, (D) DN vs CD49b, (E) DN vs DP, and (D) CD49b vs DP. Dot size is dictated by the number of genes in a given gene set and a Bonferroni-adjusted p <0.05 was used as a cut off for significantly differentially expressed pathways. Volcano plots were generated using ggplot2 in R studio with comparisons between (G) DN vs CD49b, (H) DN v DP, and (I) CD49b vs DP. Coloured dots indicate significantly differentially expressed genes based on the above inclusion criteria. N=4 biological replicates were selected from two independent experiments for RNA sequencing.

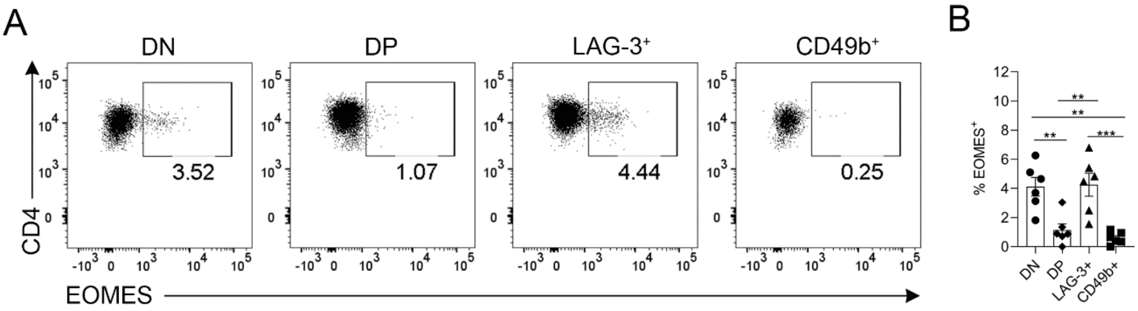


**Figure S8: Validation of EOMES.**

Dual-reporter mice were infected with X-31 IAV-i.n and on day 7 post-infection lungs were harvested and Tr1 cell populations were FACS-sorted based on LAG-3 and CD49b expression before intra-nuclear staining for EOMES. (A) Concatenated flow cytometry of EOMES expression by Tr1 cell populations. (B) Frequency of $EOMES^{+}$ Tr1 cells. Each symbol is a different biological replicate, data shown as mean +/- SEM, n=6, biological replicates total from 2 independent experiments. Statistical analysis using one-way ANOVA with Bonferroni's post-test where *p<0.05, **p<0.01, ***p <0.001.

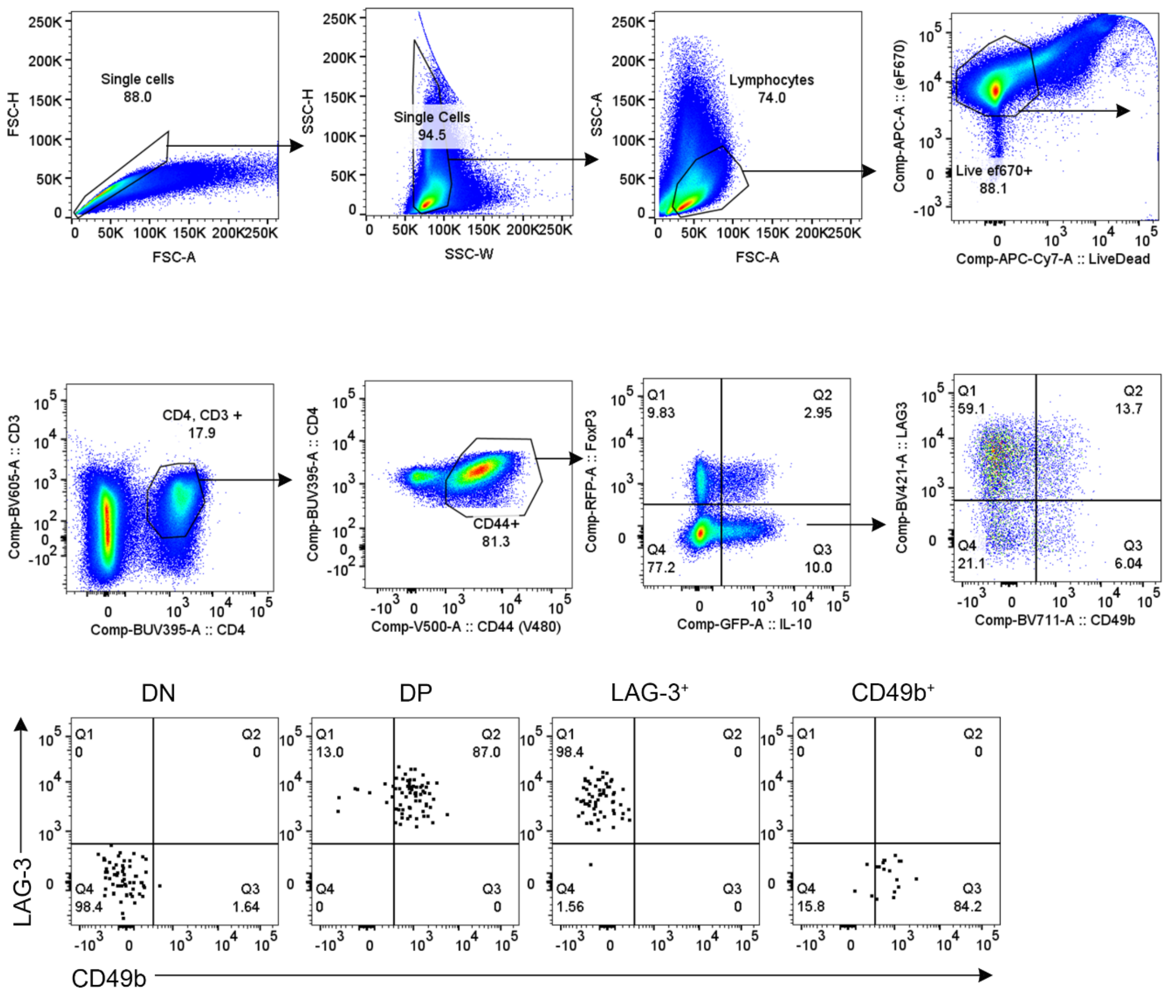


**Figure S9: Plasticity figure sort purity checks and controls.**
Dual-reporter mice were infected with X-31 IAV. Representative flow cytometry gating strategy for FACS-sorting live, CD3$^+$, CD4$^+$, CD44$^+$, FOXP3$^-$, IL-10$^+$ cells based on LAG-3 and CD49b co-expression. Representative plots showing purity of the populations post-sort are shown.

**Supplementary Table 1: Primers for RT-qPCR**

| Gene | Forward Primer (5'to 3') | Reverse Primer (5'to 3') |
|---|---|---|
| *Il10* | AAGCTCCAAGACCAAGGTGTCT | TTCTATGCAGTTGATGAAGATGTCAA |
| *Tgfb* | GAGGTCACCCGCGTGCTA | TGTGTGAGATGTCTTTGGTTTTCTC |
| *Il17a* | CCTCACACGAGGCACAAGTG | TCTCCCTGGACTCATGTTTGC |
| *Il21* | CTATGAAAATGACTTGGATCCTGAAC | CATGCTCACAGTGCCCCTTT |
| *Il22* | ACTTTCCTGACCAAACTCAGCAA | TGGTCGTCACCGCTGATG |
| *Ifng* | ATGAACGCTACACACTGCATC | CCATCCTTTTGCCAGTTCCTC |
| *Gzmb* | GACAAAGGCAGGGGAGATCAT | CGAATAAGGAAGCCCCCACA |
| *Csf1* | CTCTAGCCGAGGCCATGTG | GCTCCTCCACTTCCACTTGT |
| *Prf1* | AATATCAATAACGACTGGCGTGT | CATGTTTGCCTCTGGCCTA |

**Supplementary Table 2: Antibodies for flow cytometry**

| Antigen and Fluorophore | Clone | Cat # | Company | Final Concentration |
|---|---|---|---|---|
| 7AAD | - | 00-6993-42 | eBioscience | 5 µL /test |
| Annexin V-V450 | - | 48-8006-69 | eBioscience | 5 µL (0.025 µg)/test |
| CD25-BV421 | PC61 | 101923 | Biolegend | 1 µg/mL |
| CD3-Biotin | 145-2C11 | 100304 | eBioscience | 1 µg/mL |
| CD3-BV421 | 145-2C11 | 564008 | BD | 1 µg/mL |
| CD3-BV605 | 145-2C11 | 563004 | BD | 1 µg/mL |
| CD3-BUV805 | 145-2C11 | 749276 | BD | 1 µg/mL |
| CD3-Purified | 145-2C11 | BP0001-1 | BioXcell | 10 µg/mL |
| CD4-BUV395 | GK1.5 | 740209 | BD | 0.67 µg/mL |
| CD4-BV421 | GK1.5 | 562891 | BD | 0.67µg/mL |
| CD4-BV786 | RM4-5 | 563727 | BD | 0.67 µg/mL |
| CD8α-BUV395 | 53-6.7 | 563786 | BD | 0.67 µg/mL |
| CD8α-BUV805 | 53-6.7 | 612898 | BD | 0.67 µg/mL |
| CD44-BV480 | IM7 | 566200 | BD | 0.33 µg/mL |
| CD44-BV711 | IM7 | 563971 | BD | 0.33 µg/mL |
| CD44-FITC | IM7 | 553133 | BD | 0.33 µg/mL |
| CD44-BUV737 | IM7 | 564392 | BD | 0.33 µg/mL |
| CD49b-BV480 | HMα2 | 746355 | BD | 1 µg/mL |
| CD49b-BV711 | HMα2 | 740704 | BD | 1 µg/mL |
| CD49b-BV786 | HMα2 | 740895 | BD | 1 µg/mL |
| EOMES-PE-Cy7 | Dan11mag | 25-4875-82 | Invitrogen | 1.67 µg/mL |
| FOXP3-AF488 | MF14 | 126406 | Biolegend | 1.11 µg/mL |
| FOXP3-AF647 | MF23 | 560401 | BD | 1.11 µg/mL |
| IFNγ-PE | XMIG1.2 | 554412 | BD | 1.11µg/mL |
| IFNγ-PE-Cy7 | XMIG1.2 | 557649 | BD | 1.11µg/mL |
| IFNγ-BV480 | XMIG1.2 | 566097 | BD | 1.11µg/mL |
| IL-10-BV421 | JES5-16E3 | 566228 | Biolegend | 1.11 µg/mL |
| IL-10-PE-Cy7 | JES5-16E3 | 505026 | Biolegend | 1.11 µg/mL |
| IL-10Ra-Purified | 1B1.3A | BP0050 | BioXcell | 50 µg/mL |
| IL-17A-BV711 | TC11-18H10.1 | 506941 | Biolegend | 1.11µg/mL |
| IL-27Ra-PE | 2918 | 564337 | BD | 0.67 µg/mL |
| Ki67-BV450 | SolA15 | 48-5698-80 | Invitrogen | 1.11µg/mL |
| LAG-3-APC | C9B7W | 125221 | Biolegend | 0.8 µg/mL |
| LAG-3-BV421 | C9B7W | 125210 | Biolegend | 0.8 µg/mL |
| TCR-β –BV421 | H57-597 | 109230 | Biolegend | 0.67 µg/mL |
| TCR-β –FITC | H57-597 | 11-5961-85 | eBioscience | 0.67 µg/mL |
| TIM-3-BV421 | 5D12/TIM-3 | 747626 | BD | 0.67 µg/mL |
| TOX-PE | TXRX10 | 12-6502-82 | Invitrogen | 1.11 µg/mL |

| | | | | |
|---|---|---|---|---|
| Streptavidin BV421 | - | 563259 | BD | 0.5 µg/mL |
